# Synaptic vesicle endocytosis deficits underlie GBA-linked cognitive dysfunction in Parkinson’s disease and Dementia with Lewy bodies

**DOI:** 10.1101/2024.10.23.619548

**Authors:** D J Vidyadhara, David Bäckström, Risha Chakraborty, Jiapeng Ruan, Jae-Min Park, Pramod K. Mistry, Sreeganga. S. Chandra

## Abstract

*GBA* is the major risk gene for Parkinson’s disease (PD) and Dementia with Lewy Bodies (DLB), two common α-synucleinopathies with cognitive deficits. We investigated the role of mutant *GBA* in cognitive decline by utilizing Gba (L444P) mutant, SNCA transgenic (tg), and Gba-SNCA double mutant mice. Notably, Gba mutant mice showed early cognitive deficits but lacked PD-like motor deficits or α-synuclein pathology. Conversely, SNCA tg mice displayed age-related motor deficits, without cognitive abnormalities. Gba-SNCA mice exhibited both cognitive decline and exacerbated motor deficits, accompanied by greater cortical phospho-α-synuclein pathology, especially in layer 5 neurons. Single-nucleus RNA sequencing of the cortex uncovered synaptic vesicle (SV) endocytosis defects in excitatory neurons of Gba mutant and Gba-SNCA mice, via robust downregulation of genes regulating SV cycle and synapse assembly. Immunohistochemistry and electron microscopy validated these findings. Our results indicate that Gba mutations, while exacerbating pre-existing α-synuclein aggregation and PD-like motor deficits, contribute to cognitive deficits through α-synuclein-independent mechanisms, involving dysfunction in SV endocytosis.

## Introduction

*GBA* is the major risk gene for Parkinson’s disease (PD) and Dementia with Lewy Bodies (DLB)^1–6^, two late-onset neurodegenerative diseases, characterized by the neuronal accumulation of Lewy bodies composed of α-synuclein^7^. PD is classified as a movement disorder, although dementia affects around 50% of PD patients within 10 years after symptom onset^7^. DLB is a dementia, in which cognitive decline is generally the first and most predominant symptom^7^. Significantly, PD patients with *GBA* mutations exhibit greater and faster cognitive decline than idiopathic PD^1,7–9^. Cognitive dysfunction in both PD and DLB, which entails visuospatial, memory, and executive dysfunction, is strongly correlated with neocortical Lewy pathology^7^. However, the mechanisms through which *GBA* predisposes to cognitive dysfunction, as well as to developing α-synucleinopathies in general, are not well understood.

Homozygous or biallelic mutations in *GBA* cause the lysosomal storage disorder, Gaucher disease (GD)^10,11^. Two prevalent mutations, due to founder effects, are N370S and L444P^12,13^. GD patients have a 20-fold increased risk of developing PD accompanied by exacerbated cognitive decline. Heterozygous carriers of *GBA* mutation are at 5-fold increased risk for developing both PD and cognitive dysfunction^1,2,8,9,14–18^. In the case of DLB, *GBA* and *SNCA*, the gene for α-synuclein, are top GWAS hits. Interestingly, *GBA* mutations confer an even higher risk of developing DLB^4–6^. *GBA* encodes glucocerebrosidase 1 (GCase1), a lysosomal hydrolase responsible for breaking down the bioactive lipid glucosylceramide (GlcCer) to glucose and ceramide. In the absence of Gcase1, GlcCer and other glycosphingolipids accumulate. Interestingly, GCase1 deficiency and glycosphingolipid accumulation are also observed in post-mortem brains of patients with sporadic PD and in aging brains^19–22^. Glycosphingolipid accumulation correlates with a higher burden of α-synuclein or Lewy pathology in several brain areas^20,22,23^. These genetic, clinical, and epidemiological studies emphasize the importance of understanding mechanisms of *GBA*-linked cognitive dysfunction.

The prevailing hypothesis in the field is that *GBA* mutations lead to GCase1 deficiency, which, through a combination of lysosomal dysfunction and glycosphingolipid accumulation, trigger α- synuclein aggregation, resulting in Lewy body formation and consequently, disease associated phenotypes. We and others have shown that glycosphingolipids can directly interact with α- synuclein and promote aggregation *in vitro*^24,25^. Our long-lived mouse models of GD carrying the Gba N370S and L444P mutations, exhibited reduced GCase1 activity and accumulation of glycosphingolipids in the liver, spleen, and brain^26^. As GD patients with the L444P mutation have pronounced cognitive deficits^27^, in this study, we conducted a thorough, longitudinal examination of Gba L444P mice, in conjunction with the well-established SNCA tg PD mice that overexpress mutant human α-synuclein, and their crossbreeds, i.e. Gba-SNCA mice. We found that cognitive and motor deficits are dissociable by genotype, with Gba mutants exhibiting cognitive deficits only, SNCA transgenics motor difficulties only, and Gba-SNCA severe motor and cognitive phenotypes. Through histopathological analyses, we show that Gba L444P mutant mice lack Ser- 129-phospho-α-synuclein (pSer129α-syn) pathology but presence of the Gba mutation in Gba- SNCA mice significantly exacerbates this pathology, especially in deep layers of the cortex. Thus, cognitive deficits related to Gba mutations may emerge independently of pSer129α-syn pathology. Cortical single-nucleus RNA sequencing (snRNA-seq) analyses revealed a potential role for synaptic dysfunction, in particular synaptic vesicle endocytosis (SVE) deficits in excitatory neurons, contributing to the cognitive decline observed in Gba mutant and Gba-SNCA mice. SVE deficits are emerging as a central mechanism of PD pathogenesis, especially in rare monogenic forms of PD^28,29^. We suggest that synaptic endocytosis dysfunction also plays a role in cognitive deficits of GBA-linked PD and DLB, while greater α-synuclein pathology burden contributes to worsened motor deficits.

## Results

### Gba mutation leads to cognitive dysfunction and exacerbates motor deficits in SNCA tg mice

To determine the relative contributions of *GBA* and *SNCA* to motor and cognitive domains, we performed behavioral analyses of wild-type (WT), Gba, SNCA tg, and Gba-SNCA mouse sex-balanced cohorts. We conducted longitudinal evaluations of motor behavior every 3 months to establish the age of onset and progression of PD-like motor deficits compared to cognitive behavior deficits.

Four distinct, complementary assays were used to phenotype motor deficits: the balance beam, grip strength, hind limb clasping, and open-field locomotion tests. In the balance beam test, mice were required to walk along a narrow beam from a well-lit area to a dark, secure box^29^. The number of runs completed in a minute and the average time per run were used to assess motor performance (Fig. 1A, Supp. Fig. 1A, 1D-E). Gba mice consistently performed well on this task, comparable to WT mice, up to 12 months. In contrast, SNCA tg mice could perform this task at 3 months but began showing deficits at 6 months, which worsened by 12 months. Notably, Gba- SNCA mice demonstrated exacerbated balance deficits compared to SNCA tg mice, with significant deficits appearing as early as 6 months. By 9-12 months, Gba-SNCA mice were severely affected and unable to navigate the balance beam (Fig. 1A, Supp. Fig. 1A, 1D-E). We noted similar deficits in grip strength, measured as the force exerted by either all limbs or forelimbs of the mouse when gripping a pull bar of a grip strength meter. WT and Gba mice did not show deficits, while SNCA tg mice developed age-related decline in grip strength, which was exacerbated in Gba-SNCA mice (Fig. 1B, Supp. Fig. 1B, Supp. Fig. 2F-G).

**Figure 1:**
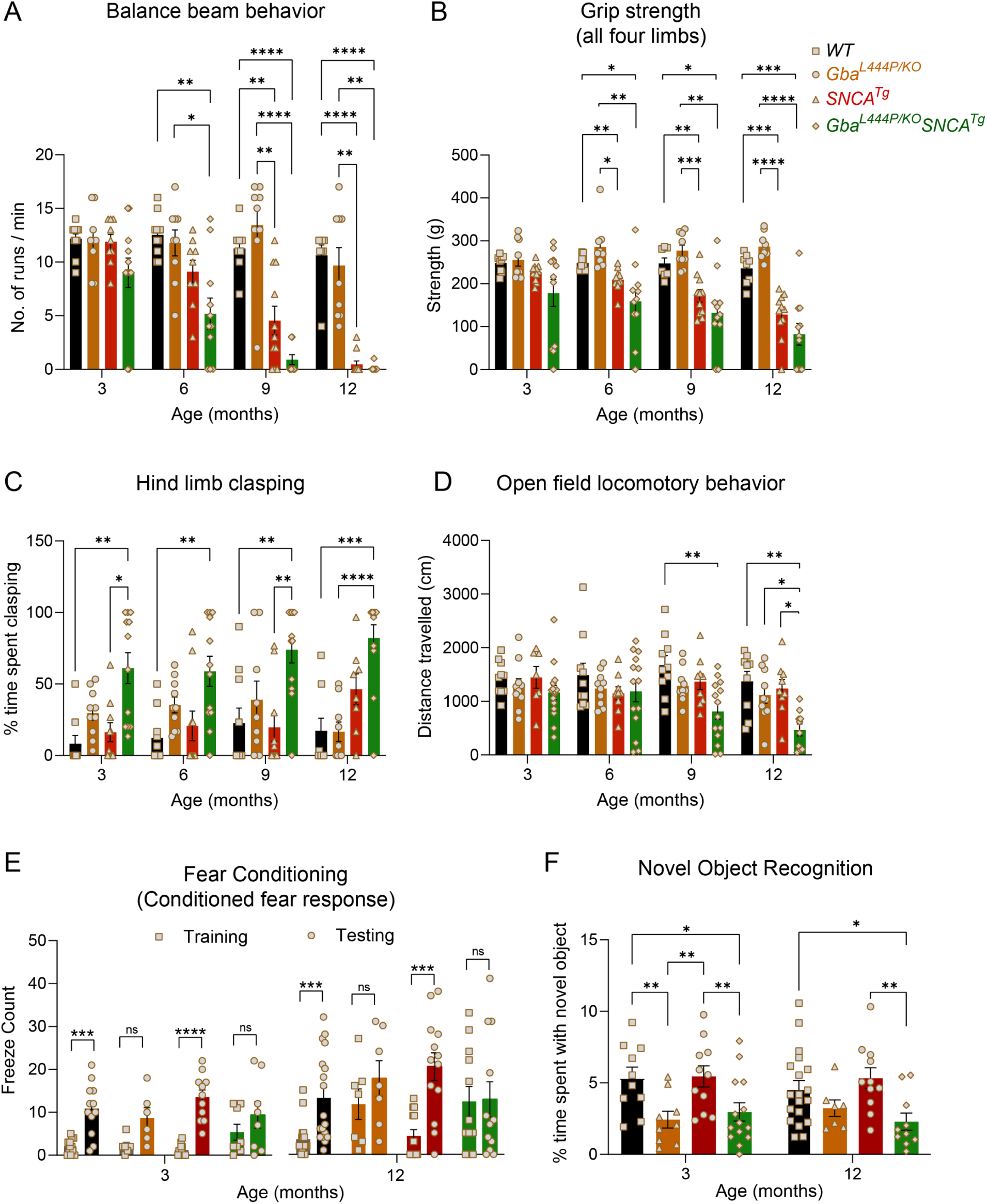
Gba mutation worsens motor deficits in SNCA tg mice and independently leads to cognitive dysfunction. WT, Gba, SNCA tg and Gba-SNCA mice cohorts were evaluated in four motor and two cognitive behavior tests in a longitudinal manner. **A.** Balance beam behavior. **B.** Grip strength of all four limbs. **C.** Hind limb clasping behavior. **D.** Open field locomotory behavior. **E.** Fear conditioning test. **F**. Novel object recognition. n = 9-12 mice for motor behavior and 6-19 for cognitive behavior, both sexes were used. Data are presented as mean ± SEM. ns - not significant; ^∗^p < 0.05, ^∗∗^p < 0.01, ^∗∗∗^p < 0.001, ^∗∗∗∗^p < 0.0001

WT Mice, when picked up by the tail and lowered towards a surface, extend their limbs reflexively in anticipation of contact. However, mice with neurological conditions display hind limb clasping. We tested the mice for 30 seconds on this maneuver and quantitated the time spent clasping (Fig. 1C). WT and SNCA tg mice did not show hind limb clasping. Gba mice showed a trend towards increased clasping (Fig. 1C, Supp. Fig. 2H), whereas Gba-SNCA mice showed a significant increase across all ages, indicating a synthetic motor phenotype (Fig. 1C, Supp. Fig. 2H).

Open field assay was performed to evaluate overall locomotion. Distance traveled exploring an open box (Fig. 1D) was comparable among all groups across ages, except for Gba-SNCA mice. At 9 months, a significant loss in exploration/locomotion was noted in Gba-SNCA mice, which worsened at 12 months (Fig. 1D, Supp. Fig. 2I). No genotypes showed anxiety-like behavior, as evaluated by the time spent in the inner/outer circles of the open field (Supp. Fig. 1C). There was no difference across the mice strains for body weight, however, Gba-SNCA mice stopped gaining weight after 6 months (Supp. Fig. 1J). In summary, motor assessments demonstrate that Gba mutants do not exhibit any appreciable motor deficits. Nonetheless, Gba mutation significantly exacerbates existing age-related motor deficits in SNCA tg mice.

Next, we evaluated the impact of Gba and SNCA mutations on cognition by employing fear conditioning and novel object recognition (NOR) tests. To avoid confounds due to learning, we performed these on two separate sets of mice at 3 months and 12 months, prior to and after the onset of motor deficits in SNCA tg mice (Fig. 1A-D). For fear conditioning, we habituated mice to standard operant boxes, followed by exposure to paired neutral stimulus (a tone) and aversive stimulus (an electric shock) on the training day. Cognitively normal mice associate this pairing and exhibit a conditioned fear response i.e. freezing when exposed to the tone alone on the testing day (24 hours later). We counted the number of freeze episodes after start of the tone and observed a conditioned fear response in WT and SNCA tg mice at both 3 and 12 months (Fig. 1E). However, Gba and Gba-SNCA mice did not show a significant response on the testing day, especially at 12 months. While this is suggestive of cognitive impairment, the results were confounded by the heightened freezing response shown by Gba and Gba-SNCA mice for aversive stimulus on the training day (Fig. 1E).

To substantiate the cognitive findings from fear conditioning, we performed a NOR test to assess recognition memory. Here, mice were presented with two similar objects (familiarization session). After 18-20 hours, one of the objects was replaced by a novel object. Mice spend more time with the novel object when cognitively normal (Fig. 1F). Gba mice spent significantly less time with the novel object at 3 months and maintained this behavior at 12 months, suggesting an early cognitive impairment (Fig. 1E). As Gba mice do not have motor problems, these results reflect memory impairments. Interestingly, SNCA tg mice did not show deficits in the NOR test (Fig. 1F), whereas Gba-SNCA do, which was comparable to Gba mice (Fig. 1F). Thus, our cognitive behavior assays suggest that the Gba mutation alone can cause cognitive impairment, with the poor cognition in Gba-SNCA mice likely driven by Gba mutation.

### Gba mutation exacerbates cortical α-synuclein pathology of SNCA tg mice

α-Synuclein aggregates, redistributes from presynaptic termini to the soma, and is phosphorylated at Ser129 in Lewy body pathology. To investigate whether Gba-mediated cognitive deficits and accelerated motor deficits are associated with α-synuclein pathology, we performed immunohistochemistry on 3- and 12-month-old mice brains, staining for α-synuclein, pSer129α-syn, and the neuronal marker NeuN. We found that Gba mice did not exhibit increased α-synuclein levels, redistribution, or accumulation of pathological pSer129α-syn in the cortex both at 3 and 12 months (Fig. 2A-C). Similar observations were made in the CA1 hippocampus and by Western blotting of whole brain homogenates (Supp. Fig. 2B-I). These findings suggest that the Gba mutation alone is insufficient to cause widespread α-synuclein pathology.

**Figure 2:**
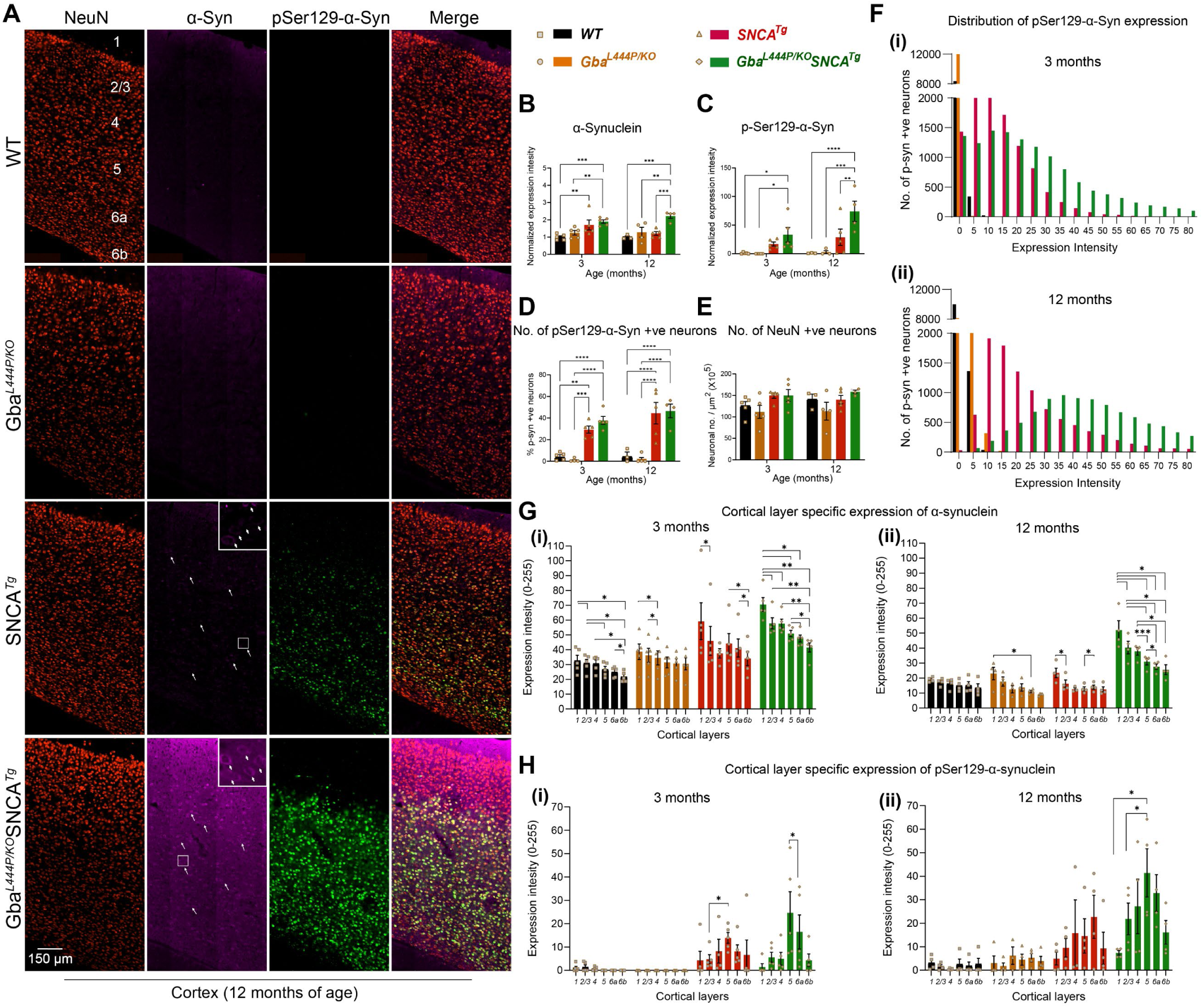
Gba mutation exacerbates α-synuclein pathology in the cortices of SNCA tg mice. **A.** Representative images of cortices of WT, Gba, SNCA tg, and Gba-SNCA mice (12 months) immunostained for NeuN (red), α-synuclein (magenta) and pSer129-α-syn (green). Note increased α-synuclein levels along with redistribution to neuronal soma (arrow) and pSer129-α- syn expression in the Gba-SNCA double mutant when compared to SNCA tg mice. **B.** Cortical α- synuclein expression at 3 and 12 months, normalized to respective WT average at each time point. **C.** Cortical pSer129-α-syn expression at 3 and 12 months, normalized to respective WT average at each time point. **D.** Percentage of cortical neurons positive for pSer129-α-syn at 3 and 12 months. **E.** NeuN positive cortical neuronal number at 3 and 12 months. **F.** Distribution of pSer129-α-syn expression intensity (Range of 0-255) in the cortical neurons at 3 months (i, WT: Mean = 0.35, 25% Percentile = 0.0024, 75% Percentile = 0.1252; Gba: Mean = 0.04, 25% Percentile = 0, 75% Percentile = 0.0058; SNCA tg: Mean = 13.05, 25% Percentile = 5.211, 75% Percentile = 18.534; Gba-SNCA: Mean = 23.781, 25% Percentile = 9.221, 75% Percentile = 33.972) and 12 months (ii), WT: Mean = 1.28, 25% Percentile = 0.439, 75% Percentile = 1.731; Gba: Mean = 1.482, 25% Percentile = 0, 75% Percentile = 2.124; SNCA tg: Mean = 26.282, 25% Percentile = 12.357, 75% Percentile = 32.834; Gba-SNCA: Mean = 53.741, 25% Percentile = 32.233, 75% Percentile = 66.691) of age. **G.** Cortical layer specific expression of α-synuclein at 3 (i) and 12 (ii) months of age. **H.** Cortical layer specific expression of pSer129 α-syn at 3 (i) and 12 (ii) months of age. n= 3-6, sex balanced. Data are presented as mean ± SEM. ns - not significant; ^∗^p < 0.05, ^∗∗^p < 0.01, ^∗∗∗^p < 0.001, ^∗∗∗∗^p < 0.0001. Scale = 150 µm

In Gba-SNCA mice, where GCase1 deficiency coexists with a pre-existing α-synuclein pathology^26^, there is significantly increased cortical α-synuclein and pSer129α-syn levels, especially at 12 months of age when compared to SNCA tg (Fig. 2A-C, Supp. Fig. 2G, I). We also noted increased redistribution of α-synuclein to the neuronal soma of Gba-SNCA mice as an independent measure of α-synuclein pathology (Fig. 2A, arrows, enlarged inserts). Interestingly, in CA1 hippocampus, the expression of α-synuclein and pSer129α-synuclein, and the soma redistribution of α-synuclein in Gba-SNCA mice were comparable to SNCA tg mice (Supp. Fig. 2B-F). Thus, in contrast to the cortex, Gba mutation only nominally exacerbates pSer129α-syn pathology, mostly in synaptic layer of CA1 at 12 months (Supp. Fig. 2F). This might be in part due to high expression of the human SNCA tg in the hippocampus^26,30^.

Next, we examined the intensity distribution of pSer129α-syn in cortical neurons as well as cortical layer specific α-synuclein and pSer129α-syn expression, at 3 and 12 months of age (Fig. 2A, F-H). Gba mice did not show pSer129α-syn pathology. Gba-SNCA mice had a higher intensity of pSer129α-syn pathology in cortical neurons at 3 months (median value of 20 vs 0 in WT and Gba, and 11 in SNCA tg mice), which further worsened at 12 months (Fig. 2F, i and ii) (median value of 47 vs 1 in WT, 0.2 in Gba, and 20 in SNCA tg mice). As the percentage of neurons expressing pSer129α-syn in Gba-SNCA was comparable to SNCA tg mice (Fig. 2D), these data suggest that the burden of pSer129α-syn per neuron in Gba-SNCA mice was greater. We did not see cortical neuronal loss in any of the mice (Fig. 2A, E), indicating that the observed behavioral deficits are not due to gross neurodegeneration.

Layer-specific analysis of α-synuclein expression revealed that cortical layer 1, which is heavily innervated by neurites, showed higher expression of α-synuclein compared to other layers. (Fig. 2G, i and ii, 3 and 12 months). Conversely, cortical layers 5 and 6a, which predominantly consist of excitatory neurons, showed higher pathological pSer129α-syn expression in SNCA tg and Gba- SNCA mice (Fig. 2H, i and ii, at 3 and 12 months), which is more evident when pSer129α-syn expression was normalized to α-synuclein (Supp. Fig. 2A, i and ii, 3 and 12 months). Together, these results suggest that the Gba mutation did not independently cause α-synuclein pathology but worsened pre-existing α-synuclein pathology in the cortex of SNCA tg mice, preferentially in layers 5 and 6a. With our behavior experiments (Fig. 1), these observations suggest that the cognitive deficits in Gba mutants emerge independently of pSer129α-syn pathology.

### Gba and SNCA driven cortical single nuclei gene expression changes

To understand cellular diversity and mechanisms for GBA-linked cognitive dysfunction, we performed single nucleus RNA sequencing (snRNA-seq) on cortical tissue from mice of all four genotypes (n=14; 3-4 mice/genotype). We chose to perform this analysis on 12-month-old mice, as Gba-SNCA mice show enhanced behavioral deficits and α-synuclein pathology, while lacking gross neurodegeneration in the cortex, allowing us to investigate disease-relevant mechanisms.

We dissected cortices and utilized our previous mouse brain nuclei isolation protocol^31^, followed by snRNA-seq on the 10X Chromium platform. We isolated 104,750 nuclei, that after cross-sample alignment, and clustering exhibited a spatial UMAP grouping uncorrelated with individual samples or genotype (Fig. 3A-E, Supp. Fig. 3A, B).

**Figure 3:**
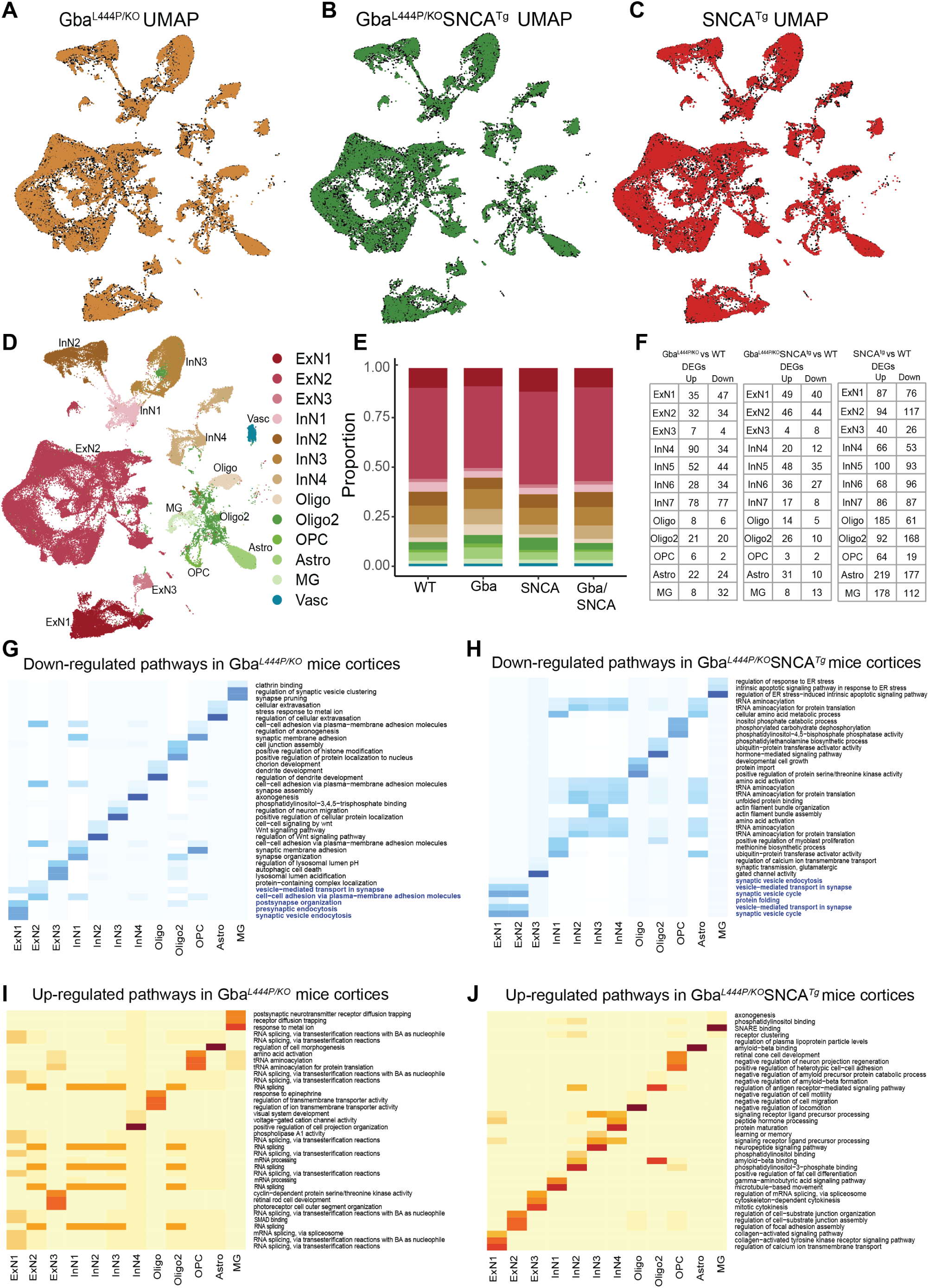
Cell type distribution, differential gene expression, and cellular pathway changes in cortex. Uniform Manifold Approximation and Projection (UMAP) dimension reduction for **A.** Gba (Amber), **B.** Gba-SNCA (Green) and **C**. SNCA tg (Red) overlayed over WT (Black) mouse cortical snRNAseq expression. **D.** UMAP showing clusters of cortical cell types identified by expression signatures. In **A-D**, UMAP 1 is shown on the x-axis and UMAP 2 on the y-axis. **E.** Proportions of the cell types in the cortices of wild type and transgenic mice. **F.** The number of differentially expressed genes (DEGs) per cell type in Gba, Gba-SNCA, and SNCA mutant mice after Bonferroni-correction for genome-wide comparisons and filtering out of genes with log2FC <I0.2I. **G-J.** Heatmap with the significantly down- and up-regulated gene ontology (GO) biological pathway alterations in 12 month old **(G, I)** Gba- and **(H, J)** Gba-SNCA mice, for each neuronal cluster type, revealed by unbiased analysis of enrichment of genome-wide corrected DEGs.

The transcriptional signatures from 104,750 nuclei, segregated into 13 broad cortical cell type clusters (Fig. 3D, E), exhibiting specific expressions of established cell-type markers (Supp. Fig. 3C). We identified three types of excitatory neurons (ExN: ExN1, ExN2, ExN3), four types of inhibitory neurons (InN: InN1, InN2, InN3, InN4), two types of oligodendrocytes (Oligo and Oligo2), oligodendrocyte precursor cells (OPC), astrocytes (Astro), and microglia (MG) (Fig. 3D). Vascular endothelial cells (Vasc) were also identified but were not further studied. The characteristic marker gene expression for each cell cluster is shown in Supp. Fig. 3C-G. Expression in all ExNs is consistent with pyramidal neurons. In ExN1, differential expression was consistent with large layer 5 pyramidal neurons shown e.g. by Fezf2 expression (Fig. 3D, Supp. Fig. 3D, F-G)). In the largest ExN subcluster, ExN2, differential expression was consistent with pyramidal neurons from several neocortical layers. The InN subclusters collectively express several classical InN markers, such as *Vip*, *Sst*, *Erbb4*, while the subclusters InN1 and InN3 specifically contain layer 2/3 interneurons (Supp. Fig. 3C). Typical marker signatures used for Oligo and Oligo2 (*Mbp*, *Ptgds*, *Mal*), suggests that Oligo2 also contains minor neuronal populations in addition to oligodendrocytes (Supp. Fig. 3C). The OPC (*Vcan*, *Epn2*, *Tnr*), Astro (*Aqp4*, *Prex2*, *Luzp2*) and MG (*Cd74*, *C1qa*, *Csf1r*) markers are consistent with prior literature^31,32^. The relative proportions of major cell types were roughly similar between the four genotypes (Fig. 3E).

Next, we examined the expression of endogenous Gba, mouse Snca in WT brains and the expression of human transgenic SNCA (hSNCA) using Thy-1 in SNCA tg brains (Supp. Fig. 3H- J). Snca is enriched in excitatory neuronal clusters, including ExN1, consistent with published literature (Supp. Fig. 3H)^33,34^. This pattern was also true for hSNCA^34–36^, although hSNCA is also expressed in glial cell types in SNCA tg mice (Supp. Fig. 3H, I). The high hSNCA expression in the predominantly layer 5-populated ExN1 cluster (Supp. Fig. 3H) is consistent with the layer 5 specific increases of α-synuclein pathology demonstrated by immunohistochemistry (Fig. 2H, Supp. Fig. 2A). In contrast, Gba is generally found at low levels in brain cells^37–39^ (Supp. Fig. 3J, i and ii).

### Mutant Gba drives transcriptional downregulation of synaptic pathways in neurons

After correcting for genome-wide comparisons, we identified up- and down-regulated differentially expressed genes (DEGs) in all cell types in Gba, Gba-SNCA and SNCA tg mice cortices compared with WT (Fig. 3F, Supp. Table 1-3). To gain insights into the cognitive deficits seen in Gba mice, we focused on DEGs in neuronal clusters, comparing Gba with WT (Supp. Table 1). Strikingly, Gba mutant mice showed a general downregulation of many genes functioning at the synapse (*Arc*, *Syp*, *Actb, Nrg1*, *Nlgn1*, *Nrg1*, *Grm7*, *Grip1*, *Ptprd*, *Nlgn1*, *Il1rapl2*, *Gabra1*, *Cntn5*, *Lingo2*, *Erbb4*, *Nptn*, *Lrrtm4*, *Actb*, *Cntnap2*, *Lrfn5*), suggestive of a synaptic dysregulation signature related to Gba. *Ahi1*, a gene important for cortical development and vesicle trafficking was upregulated in all neuronal classes in Gba mice.

To define the major pathways impacted, we performed unbiased gene ontology (GO) enrichment analysis comparing Gba to WT (Fig. 3G, I). As shown by heatmaps depicting the top biological pathway changes, we found a consistent decrease in synaptic pathways in cortical ExNs of Gba mice driven by reduced expression of *Syp*, *Actg1*, *Actb*, *Nlgn1*, *Grm8*, *Nrg1*, *Arc* (Fig. 3G). ExN1 and ExN2, shared robust downregulation of genes involved in SVE, presynaptic endocytosis, and vesicle-mediated transport in synapse in Gba mice (Fig. 3G, highlighted, Supp. Table 4). Additionally, in Gba mice, ExN1 and ExN2 showed downregulation of cellular pathways and genes involved in both pre- and postsynapse organization, and synaptic protein-containing complex localization (Fig. 3G, highlighted, Supp. Table 4). In the smaller ExN3 cluster, DEGs were fewer and involved in lysosomal lumen acidification (Supp. Table 4). In contrast, the significant upregulated pathways in ExN1 and ExN2 of Gba involve RNA splicing (Fig. 3I, Supp. Table 4). Inhibitory neurons (InN1-4) in Gba mutant mice showed downregulation of multiple synapse-associated pathways, including genes involved in synapse organization, synapse membrane adhesion in InN1, synapse assembly in InN4, Wnt signaling in InN2, and axonogenesis (Fig. 3G, Supp. Table 4). The upregulated pathways in InN1-4, similar to ExNs, are related to RNA splicing (Fig. 3I, Supp. Table 4).

Next, we compared DEGs in neuronal clusters in Gba-SNCA with WT (Supp. Table 2). Consistent with our finding of a Gba-driven synapse effect, ExN clusters in Gba-SNCA mice show robust downregulation of synapse related genes (Supp. Table 4). GO enrichment analysis comparing Gba-SNCA to WT revealed the top downregulated pathways in cortical ExN1 and ExN2 were SV cycle, vesicle-mediated transport in synapse, and SVE (driven by *Actb*, *Actg1*, *Unc13a*, *Cacna1a*, *Calm1*, *Btbd9*, *Prkcg*, *Pacsin1* reductions Fig. 3H, highlighted). Although the individual DEGs between Gba and Gba-SNCA are not identical, the synaptic pathways being impacted are highly similar, suggestive of a common synaptic dysregulation signature related to Gba (Compare Fig. 3G with 3H). In Gba-SNCA InNs we see distinct pathways such as ubiquitin-protein transferase activator activity in InN1 and tRNA aminoacylation in InN2-4 being down regulated (Fig. 3H). The upregulated pathways in Gba-SNCA in ExNs are related to focal adhesion assembly and in InNs are diverse and include amyloid binding (Fig. 3J).

Next, we analyzed all neuronal DEGs through SynGO to define synaptic DEGs in Gba and Gba- SNCA mice. The SynGO analysis revealed more significant suppression of synaptic genes in ExN classes compared to InN classes in both genotypes (Fig. 4A-D). Cnet plots revealed enrichment of converging and predominantly down-regulated synaptic genes and pathways in both Gba and Gba-SNCA mouse cortices (Fig. 4E, F), consistent GO analyses (Compare Fig. 3G, H, with Fig. 4E, F).

**Figure 4:**
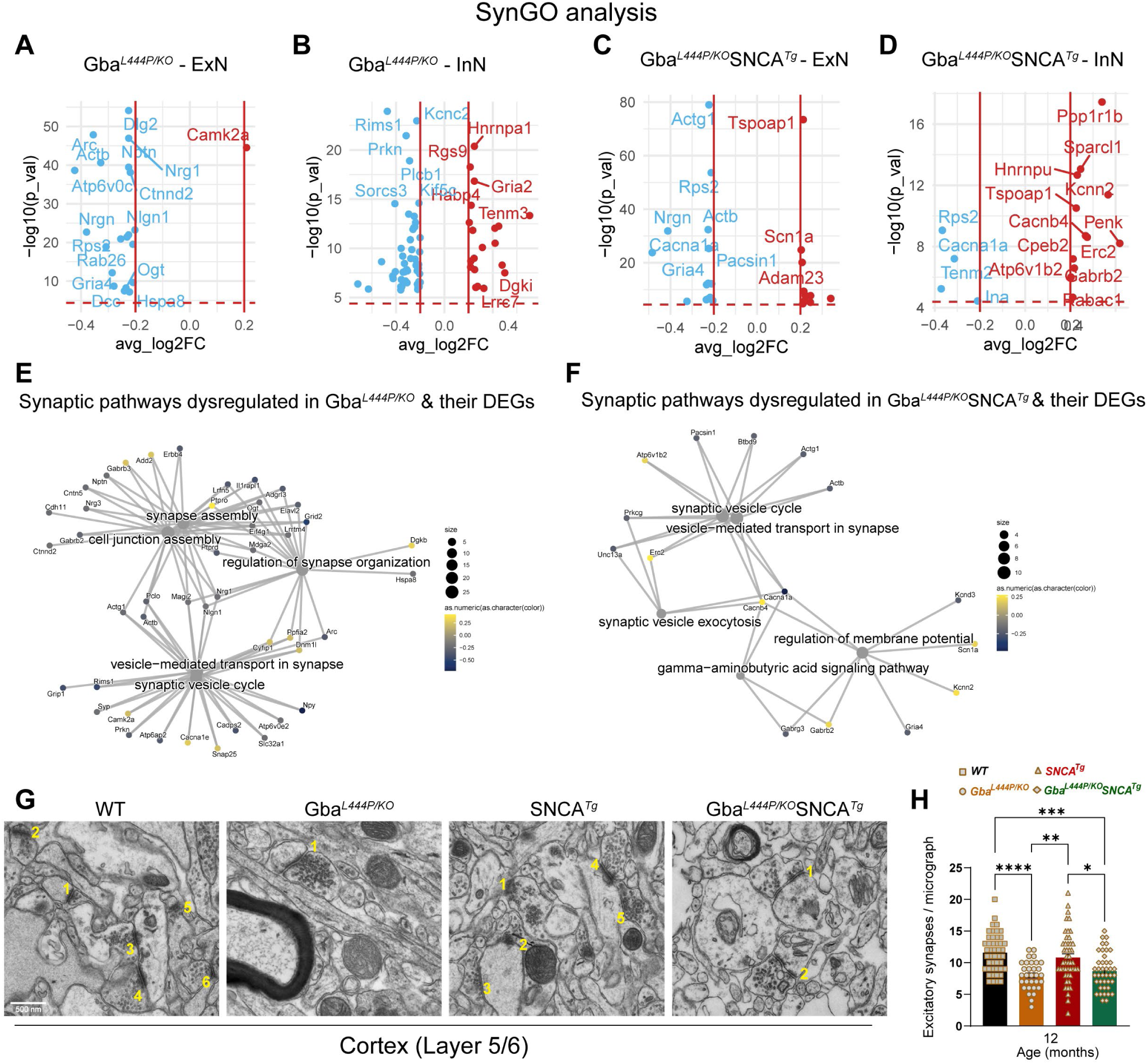
Analysis of synapse related gene expression in Gba and Gba-SNCA neurons shows similar deficits in synapse vesicle cycling. **A-D.** Analysis of significant DEGs that participate in synapse function, as annotated by SynGO, after Bonferroni-correction in excitatory (ExN1-3, **A, C**) and inhibitory (InN1-4, **B, D**) neurons. All genes with log2FC <0.2 were filtered out. **E, F.** Cnet plots of differentially expressed synapse associated genes as annotated by SynGO, in **(E)** Gba-, and **(F)** Gba-SNCA mice cortices, after Bonferroni-correction for multiple comparisons. **G.** Electron micrographs of cortical layer 5/6 number for excitatory synapses. **H**. Quantitation of excitatory synapses in the cortical layer 5/6. Data are presented as mean ± SEM, Scale = 500 nm, *p<0.05, ***p<0.001., N=2 brains in WT, SNCA tg, and Gba-SNCA mice, and 1 brain in Gba mutant mice. 23-25 micrographs/genotype.

To evaluate if the downregulation of synapse organization pathways leads to a decrease in excitatory synapse number, we performed electron microscopy on 12-month old cortex samples. Electron microscopy was chosen as it allows for accurate quantification of synapse numbers while avoiding problems of individual synaptic protein marker differences across genotypes. As seen in Fig. 4G-H, number of excitatory synapses in deep layers of cortex is indeed reduced in Gba mutant and Gba-SNCA mice compared to SNCA and WT mice.

To assess SVE changes at the protein level, we immunostained cortex and hippocampus sections, for the SVE protein, endophilin A1 (a risk allele for PD) and the endocytic lipid PIP2. Conforming with the snRNA expression data, endophilin A1 and PIP2, showed a decreased trend in the cortex (Fig. 5A, Cortex, B, C). Interestingly, in the synaptic layer in CA1 of the hippocampus where endophilin A1 and PIP2 are enriched, we noted significantly reduced endophilin A1 and PIP2 expression in Gba and Gba-SNCA mice (Fig. 5A, CA1 hippocampus, D, E). Together, these observations corroborate our findings from snRNA-seq analysis of Gba-driven suppression of SVE genes and show these deficits are not limited to the cortex.

**Figure 5:**
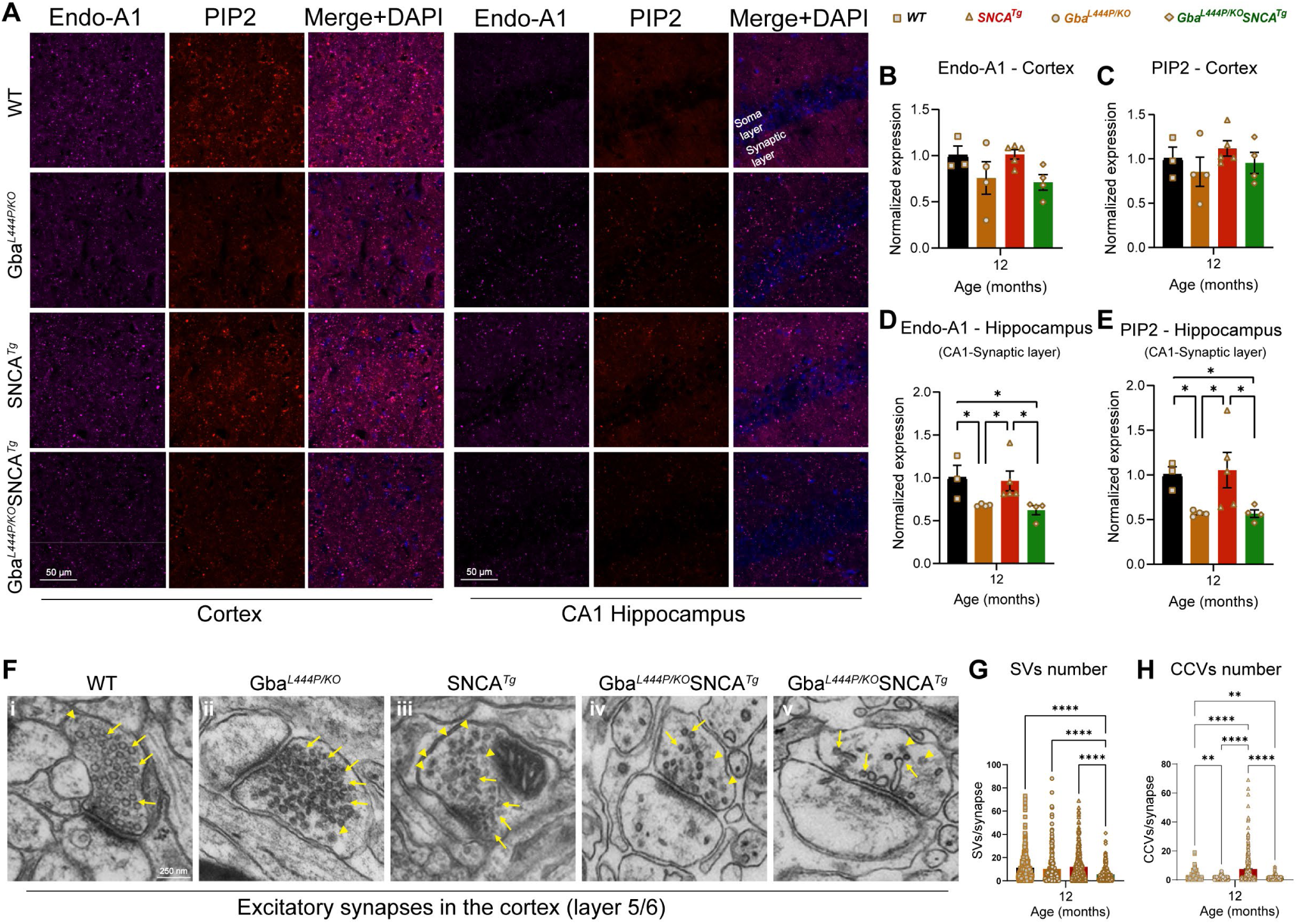
Decreased expression of SVE markers and loss of SVs in Gba mutants. **A**. Representative images showing cortical and CA1 hippocampal expression of endophilin-A1 (Endo-A1) and phosphatidylinositol 4,5-bisphosphate (PIP2), two markers of synaptic vesicle endocytosis, in WT, Gba, SNCA tg, and Gba-SNCA mice at 12 months of age. **B.** Cortical Endo- A1 expression at 12 months, normalized to WT average. **C.** Cortical PIP2 expression at 12 months, normalized to WT average. **D.** Endo-A1 expression in the CA1 Hippocampal synaptic layer, normalized to WT average. **E.** PIP2 expression in the CA1 hippocampal synaptic layer, normalized to WT average. Data are presented as mean ± SEM. Scale = 50 µm. * p<0.05. n=4-5 brains/genotype. **F.** Electron micrographs of excitatory synapses in cortical layer 5/6 showing synaptic vesicles (SVs, arrows) and clathrin-coated vesicles (CCVs, arrowheads) in WT (**i**), Gba mutant (**ii**), SNCA tg (**iii**), and Gba-SNCA (**iv** and **v**) mice. Note SVs with variable shapes and sizes in Gba-SNCA synapse **(v). G-H**. Quantitation of SVs (**G**) and CCVs (**H**) in the excitatory synapses of the cortical layer 5/6. Data are presented as mean ± SEM, Scale = 250 nm, *p<0.05, **p<0.01, ****p<0.0001, N=2 brains in WT, SNCA tg, and Gba-SNCA mice, and 1 brain in Gba mutant mice. 23-25 micrographs, 150-300 synapses, per genotype.

To determine whether these transcriptional and protein expression changes affect the SV cycle, we examined electron micrographs of excitatory synapses in the cortical layer 5/6 and quantified SVs and clathrin coated vesicles (CCVs), which serve as proxies for SV cycling and SVE (Fig. 5F-I). In the Gba mutant mice, the number of SVs was comparable to that in WT mice, however, a significant loss of CCVs was observed, potentially indicating a slowdown in clathrin-mediated SVE (Fig. 5F-H). In Gba-SNCA mice, there was a marked reduction in both SVs and CCVs (Fig. 5F-H), with several synapses showing SVs with variable shapes and sizes (Fig. 5F, v), indicating a severe disruption of SVE and SV recycling. Interestingly, the SNCA tg mice displayed a significant increase in CCVs (Fig. 5F and H), consistent with findings in other α-synuclein models^40,41^. Together, these findings reveal a distinct pattern of SVE disruption in Gba mutant mice, which is exacerbated in Gba-SNCA mice, with altered SV recycling potentially leading to cognitive dysfunction.

### ExN1 cluster contains vulnerable layer 5 cortical neurons

As cortical layer 5 neurons exhibited the highest vulnerability in terms of α-synuclein pathology, (Fig. 2A, H, Supp. Fig. 2A), and transcriptional changes associated with SVE (Fig. 3G, H), we further investigated the ExN1 cluster which has high hSNCA transgene and Gba expression (Supp. Fig. 3I, J). Upon subclustering, ExN1 was divided into six subclusters (Supp. Fig. 4A-D), all of which contained cells expressing the layer 5 marker *Fezf2* (Supp. Fig. 4D, E). One subcluster, ExN1.1, was characterized by high expression of Arc (Supp. Fig. 4D), which is downregulated in Gba mutant neurons (Supp. Fig. 4F). Our analysis confirmed the greatest downregulation of synapse-associated genes in ExN1.1 subcluster in both Gba and Gba-SNCA mice (Supp. Fig. 4D, F-I). In contrast, non-synaptic DEGs were evenly up- and downregulated (Supp. Fig. 4F, G). SV cycle pathways were consistently downregulated throughout ExN1, largely driven by the same genes identified in our non-targeted analysis (Fig. 3G, H). Additionally, Rab26, a key regulator of SV endocytosis and autophagy^42^, was downregulated in both Gba genotypes (Supp. Fig. 4H, I). These findings suggest that layer 5 excitatory neurons are selectively vulnerable because of both high hSNCA and Gba expression and SV cylcing deficits driven by Gba mutations. These mechanisms appear to act synergistically to exacerbate α-synuclein pathology in Gba-SNCA mice.

### Modest glial transcriptional changes in Gba mutant and Gba-SNCA cortex

Compared to neuronal clusters, glial clusters in Gba and Gba-SNCA cortices exhibited fewer DEGs (Fig. 3F, Supp. Table 1-3). In Gba cortex, MG showed altered gene expression patterns indicative of reduced synaptic remodeling. Notably, synapse pruning and regulation of SV clustering pathways were downregulated (Fig. 3G, Table 4). Postsynaptic neurotransmitter receptor diffusion trapping was upregulated (Fig. 3I, Supp. Table 4). These changes reinforce synapse dysfunction as a central pathological mechanism in Gba mutant cortex. The down and upregulated pathways in astrocytes in Gba are related to cellular extravasation and morphology (Fig. 3I, Supp. Table 4). There were fewer DEGs in OPCs and Oligodendrocytes (Fig. 3F); therefore, clear pathway differences are harder to discern (Fig. 3G-I).

In Gba-SNCA, MG exhibit decreased endoplasmic reticulum stress response, while SNARE binding and aspects of phosphatidylinositol binding were upregulated (Fig. 3H, J, Supp. Table 4). Gba-SNCA Astro exhibit decreased tRNA aminocylation and increased amyloid-beta binding (Fig. 3H, J). In Gba-SNCA OPCs, phosphatidylinositol binding was downregulated, while Oligodendrocytes showed modest pathway changes (Fig. 3H, J). Overall, in Gba mice, all glial cell types have muted responses, while Gba-SNCA mice had slight astrocytic activation. To confirm our analysis, we immunostained with the microglial marker Iba1, CD68 for activated microglia, and GFAP for astrocytes (Supp. Fig. 5). We did not observe any significant increase in Iba1 or CD68+ve microglial number, suggesting negligible microglial activation in Gba and Gba- SNCA cortex. We observed a trend towards increased GFAP levels in Gba-SNCA mice compared to WT (Supp. Fig. 5A-D). Together, these data suggest that glial responses are modest, consistent with snRNAseq data.

### Cortical transcriptional changes indicate broad synapse dysregulation in SNCA cortex

SNCA tg mice showed the greatest number of DEGs compared to WT relative to Gba and Gba- SNCA mice (Fig. 3F, Supp. Table 3). The neuronal clusters showed broad alterations of synapse related gene expression. Both up and down-regulated DEGs are involved in synapse assembly, regulation of synapse structure or activity, and postsynapse organization (Supp. Figs. 6A-B, Supp. Table 3). SynGO analysis of these DEGs revealed regulation of synapse structure and function as the main pathway impacted in SNCA tg mice (Supp. Fig. 6C-D). As excitatory synapse number was not changed significantly in cortical regions of SNCA tg mice (Fig. 4G, H) compared to WT, this likely results in functional deficits. Consistent with the observed cortical α-synuclein pathology at this age (Fig. 2A, D), unfolded protein handling was upregulated, as were pathways involved in protein folding and refolding specifically in ExN1 and ExN2 clusters (Supp. Fig. 6B, Supp. Table 3). Additionally, regulation of protein ubiquitination was upregulated in all ExNs (Supp. Fig. 6B, Supp. Table 3). OPCs and oligodendrocytes show changes related to oligodendrocyte differentiation. MG showed down-regulation in immune receptor binding and up-regulation in cation channel activity (Supp. Fig. 6A-B, Supp. Table 3). In Astros, cell junction assembly was decreased, and ion channel activity was increased (Supp. Fig. 6A-B). However, we did not observe significant microgliosis or astrogliosis in SNCA tg mice cortices by immunohistochemistry (Supp. Fig. 5). Despite SNCA transcriptional changes, Gba signatures are predominant in Gba- SNCA cortices.

## Discussion

Surveys of PD and DLB patients and their caregivers highlight that maintaining cognitive abilities is a major unmet need^43^. The *GBA* gene is an ideal choice to investigate this non-motor symptom, because it is the most common risk gene for PD^8^ and *GBA* mutations are linked to cognitive deficits in both diseases^8,18^. Here, we present detailed age-dependent behavioral and pathological phenotyping of the Gba-SNCA line alongside WT, Gba mutant, and SNCA tg mice. We demonstrate that Gba-SNCA mice recapitulate both cognitive dysfunction and motor deficits seen in GBA-linked PD and DLB. Significantly, we carried out snRNA-seq analysis of the cerebral cortex in these four mice genotypes and have built one of the first comprehensive *Gba* transcriptomic data sets. We encourage the scientific research community to utilize this rich dataset as a resource for additional analyses on PD and DLB (NCBI GEO GSE283187).

### Gba-SNCA as a mouse model of *GBA*-linked PD and DLB

Cognitive dysfunction has been noted in several existing mice models of PD designed to study motor deficits, including those focused on α-synuclein pathology^44–47^. These models involve overexpression of hSNCA mutations or the use of pre-formed fibrils to induce α-synuclein aggregation^48^ and have been instrumental in elucidating the mechanisms of α-synuclein pathology and its impact on neurodegeneration and cognitive decline. However, they do not fully replicate the complex genetic and pathological features of human PD and DLB, importantly, the contribution of *GBA*. Additionally, most biallelic Gba models are hampered by early lethality, precluding age-related studies, and hence investigated as heterozygotes^48,49^. Here, we build on our previous analyses of long-lived biallelic, Gba mutant mice and Gba-SNCA^26^ and show that Gba-SNCA mice are an excellent model of GBA-linked PD and DLB. Gba-SNCA mice offer significant advancements as they exhibit worsened motor deficits compared to SNCA tg in an age-dependent manner as well as early cognitive deficits, closely mirroring the human condition. This is further evidenced by exacerbation of cortical α-synuclein pathology in Gba-SNCA mice. Significantly, by comparing Gba, SNCA tg, and Gba-SNCA mice, we were able to demonstrate that the Gba mutation alone can drive cognitive dysfunction. These data are congruent with recent studies using heterozygous L444P Gba mutant mice^49,50^. Notably, Gba-SNCA mice exhibited enhanced motor deficits, but cognitive deficits were on par with Gba mice, matching the similar synaptic pathway deficits in their neuronal populations as assessed by snRNAseq. In sum, Gba-SNCA mice capture the complexities of *GBA*-linked PD and DLB and serve as a good mouse model for these synucleinopathies.

### GBA-linked cognitive dysfunction is independent of α-synuclein pathology

A striking finding is that cognitive dysfunction occurs independent of or precede α-synuclein pathology in Gba mutants. pSer129α-syn is the gold standard to define Lewy bodies in both PD and DLB^51–57^ ^56,58,59^. Yet, we did not detect any pSer129α-syn in Gba mutant brain nor did we observe redistribution of α-synuclein to the soma, a posttranslational modification independent measure of pathology^60^. This was most evident in the hippocampus, where the synaptic and cell body layers are demarcated. While α-synuclein pathology in cortical areas does lead to cognitive dysfunction in mice overexpressing mutant α-synuclein or those injected with PFFs^47^, our study specifically challenges the necessity of α-synuclein pathology in the development of cognitive deficits in *GBA* mutations. Clinical support for this comes from children with neuropathic forms of Gaucher disease, who show cognitive deficits but do not develop neocortical α-synuclein pathology^61,62^. While a recent study has made similar observations focusing on the hippocampus^50^, it would be interesting to explore whether aging Gba mice could initiate α-synucleinopathy or if other pathological forms of α-synuclein are involved.

In contrast to cognitive deficits, motor deficits are strongly related to α-synuclein pathology in Gba- SNCA mice. Notably, we observed significant α-synuclein pathology in cortical layers 5, similar to that seen in other PD mice models and human PD and DLB patients^34,63^. Our snRNA-seq data suggest that layer 5 ExNs are also vulnerable to Gba mediated synaptic dysfunction, which could contribute to an increased accumulation of α-synuclein pathology in cortical layer 5, and in turn, the severity of the motor deficits. Thus, Gba-SNCA mice also highlight specific cortical neuronal vulnerabilities, allowing for further investigations into cortical mechanisms of PD and DLB.

### SVE and organization deficits in Gba-linked cognitive dysfunction

Our snRNAseq analysis showed clear evidence of specific synaptic changes. Many synaptic genes that function in SV cycle, SVE, synapse organization, synapse membrane adhesion, and synapse assembly were downregulated in neuronal clusters in the cortex of Gba as well as in Gba-SNCA mice, suggesting common synaptic dysfunction mechanisms linked to Gba. Because Gba mutant mice do not show α-synuclein pathology, this implies that synaptic dysfunction directly contributes to the observed cognitive deficits, rather than a consequence of disease pathology or neurodegeneration. In support of this tenant, we observed excitatory neuronal synapse loss in the cortex of Gba mutant and Gba-SNCA mice. In other dementias such as Alzheimer’s disease, synapse loss correlates tightly with cognitive decline^64^. Our findings suggest that this is likely true for GBA-linked PD and DLB, in line with available clinical studies^65–67^.

SVE was the major pathway downregulated in neurons in both Gba mutant and Gba-SNCA mice, supported by our immunohistochemistry and electron microscopy experiments. Three key genes driving this pathway are *Hspa8*, *Dlg2 and Arc*. *Hspa8* (encoding HSC70) contributes to synapse vesicle uncoating and functions with *Dnajc6/PARK19*, a familial PD gene^28,68^. *Dlg2* is a risk gene for sporadic PD that participates in receptor clustering in synapses^69^. *Arc* is an activity regulated gene that regulates transcription of many synaptic and SVE genes^70^. Interestingly, both clinical and experimental data link SVE deficits to cognitive deficits. The levels of dynamin1, the endocytic GTPase, correlate with Lewy Body Dementia^71^. Patients with mutations in *DNAJC6/PARK19* and *Synj1/PARK20* which encode two key SVE proteins--auxilin and synaptojanin1—have cognitive deficits. Endocytic mutant mice show deficits in NOR and fear conditioning^72,73^, supporting the tenet that SVE deficits lead to cognitive dysfunction. We suggest that *Arc* could serve as an upstream regulator of the synaptic transcriptional changes seen in Gba and Gba-SNCA cortices. Another contributor to the Gba-driven alteration of SV cycling is plasma membrane lipid composition changes. Emerging evidence indicates altered sphingolipid composition, such as in Gba mutants, may interfere with phosphoinositide biology at membranes^74,75^. Future research should address the topic of co-regulated lipids.

### Limitations

The Gba-SNCA mouse model effectively replicates both behavioral and histopathological characteristics of GBA-linked PD and DLB, offering a valuable tool for future research. Our study highlights the critical role of synaptic dysfunction in GBA-linked cognitive decline, occurring independent of α-synuclein pathology. While transcriptional analysis supported by histology revealed significant disruptions in SVE and synapse assembly in cortical excitatory neurons of both Gba and Gba-SNCA mice, the upstream regulators of these changes remain to be identified. While glial cells exhibit a more modest transcriptional response compared to neurons at this age, their involvement in synaptic dysfunction at earlier stages cannot be ruled out. Our comprehensive transcriptomic dataset provides a rich resource for further exploration, and we encourage the scientific community to leverage this data to elucidate GBA-linked disease mechanisms.

## Methods

### Mice

Gba mutant mice have been previously described in Mistry et al. (2010)^76^ and Taguchi et al. (2017)^26^. These mice have a copy of the Gba L444P mutant allele and a Gba KO allele, with Gba expression rescued in skin to prevent early lethality. SNCA tg mice overexpress the human α-synuclein A30P transgene (heterozygous), and have also been previously described^30^. Gba mutant mice were crossed to SNCA tg to obtain Gba-SNCA double mutant mice. Age and sex matched WT mice were used controls.

### Behavior evaluation

WT, Gba, SNCA tg, and Gba-SNCA mice were examined for motor behavior longitudinally at 3, 6, 9, and 12 months of age as described previously (n=9-12 mice/genotype, sex-matched)^29^. The balance beam test assesses the ability to walk straight on a narrow beam from a brightly lit end towards a dark and safe box. Number of times a mouse could perform this behavior in a minute and the average time taken for each run were evaluated. The grip strength of all the limbs and the forelimbs was assessed by measuring the maximum force (g) exerted by the mouse in grasping specially designed pull bar assembly in tension mode, attached to a grip strength meter (Columbus Instruments, Ohio, USA). Mice, when picked up by the base of the tail and lowered to a surface, extend their limbs reflexively in anticipation of contact. Mice with certain neurological conditions display hind limb clasping instead of extension. Mice were tested on this maneuver for 30 secs and the hindlimb, forelimb, and trunk clasps were scored (0: no clasp; 1: one hind limb clasp; 2: both the hind limbs clasp; 3: Both the hind limbs clasping with at least one forelimb clasp; 4; Both the hind limbs clasp with trunk clasping). For evaluation of overall locomotory capabilities, mice were allowed to explore an open field arena for 5 minutes, which was videotaped to assess the distance travelled using Noldus Ethovision CT software.

To evaluate cognition, we employed fear conditioning and novel object recognition (NOR) tests. To avoid learning-induced confounding factors, we performed these tests on two separate sets of mice at 3 and 12 months. For fear conditioning test, we initially habituated mice in standard operant boxes for 2 minutes, followed by exposure to a 30-second neutral stimulus (a 80 dB tone), which ended with 2 seconds of an aversive stimulus (a 0.1 to 1.0 mA electric shock). This pairing associates the neutral stimulus with fear, leading the mice to exhibit fear responses, such as freezing, when exposed to the tone alone. We tested for freezing 24 hours later on the testing day. Cognitively normal mice will form a conditioned fear response, exhibiting increased freezing behavior on the testing day compared to the training day after exposure to the tone alone, indicating their ability to associate tone with the electric shock they received on the training day. We measure this conditioned fear response as the number of freeze counts after starting the tone for a total of 3 minutes. The NOR test was used to assess the recognition memory. First, mice were acclimatized to the novel object arena without any objects in it. After 24 hours, familiarization session was performed where mice were presented with two similar objects for 8 minutes. After 18-20 hours, one of the two objects was replaced by a novel object and mice were allowed to explore for 8 minutes. Mice being exploratory animals, spend more time with novel object when their cognition is normal, which we used as a measure of NOR test.

### Immunohistochemistry

Equal number of male and female mice at 3 and 12 months of age (n=3-6/genotype) were used for immunohistochemistry. Mice were anaesthetized using isoflurane inhalation and perfused intracardially with chilled 0.9 % heparinized saline followed by chilled 4 % paraformaldehyde (PFA) in 0.1 M phosphate buffer (PB). The brains were post-fixed in the same buffer for ∼48 hours and cryoprotected in increasing grades of buffered sucrose (15 and 30 %, prepared in 0.1 M PB), at 4 °C, and stored at −80 °C until sectioning. Sagittal brain sectioning (30 μm thick) was performed using a cryostat (Leica CM1850, Germany), and the sections were collected on gelatinized slides, and stored at -20 °C until further use. For immunofluorescence staining, sections were incubated in 0.5 % triton-X 100 (Tx) (15 mins), followed by incubation in 0.3 M glycine (20 mins). Blocking was performed using 3% goat serum (90 mins), followed by overnight incubation (4° C) in the primary antibodies. Following day, sections were incubated in Alexa-conjugated secondaries for 3-4 hours, followed by coverslip mounting using an antifade mounting medium with (H-1000, Vectashield) or without (H-1200, Vectashield) DAPI. Coverslips were sealed using nail polish. 1X PBS with 0.1 % Tx was used as both washing and dilution buffer. Below is the list of antibodies used and their dilutions.

| Antibody | Dilution | Manufacturer, RRID |
| --- | --- | --- |
| Mouse Anti-α-Synuclein | 1:500 | BD Biosciences (610786), AB_398107 |
| Guinea Pig Anti-NeuN | 1:500 | Sigma-Aldrich (ABN90P), AB_2341095 |
| Rabbit Anti-α-synuclein (phospho S129) | 1:800 | Abcam (ab51253), AB_869973 |
| Rabbit Anti-Iba1 | 1:300 | Wako Chemicals (019-19741), AB_839504 |
| Guinea Pig Anti-GFAP | 1:400 | Synaptic Systems (173004), AB_1064116 |
| Mouse Anti-CD68 | 1:400 | Invitrogen (MA1-80557), AB_929627 |
| Rabbit Anti-Endophilin 1 | 1:200 | Synaptic Systems (159002), AB_887757 |
| Mouse Anti-PIP2 | 1:200 | Invitrogen (MA3-500), AB_568690 |

### Fluorescence slide scanner, confocal microscopy, and image analysis

Fluorescent images were acquired using a fluorescence slide scanner (VS200, Olympus) or confocal microscope (LSM 800, Zeiss) with a 40X objective using appropriate Z-depth. Images were then analyzed using FIJI software from National Institute of Health (NIH), blinded for genotype. Whole cortex was demarcated as per Paxinos and Franklin, 2008. After performing sum intensity projection, the expression intensity was measured on an 8-bit image as the mean gray value on a scale of 0–255, where ‘0’ refers to minimum fluorescence and ‘255’ refers to maximum fluorescence. For counting NeuN+ neurons, images were thresholded using the ‘otsu’ algorithm and the cells larger than 25 mm^2^ were counted using the ‘analyze particles” function^29^. A similar method was used to count Iba1+ve microglial cells, and CD68 +ve cells. GFAP+ astroglial cells were counted manually using the ‘cell counter’ function. Regions of interest (ROIs) obtained for individual NeuN+ were overlayed on pSer129α-syn staining to obtain numbers of neurons that were positive for pSer129α-syn. To analyze the cortical layer-specific expression of α-synuclein and pSer129α- syn, we initially determined the proportions of cortical layers (1 to 6b) in sagittal brain slice images obtained from the Allen Brain Atlas (https://mouse.brain-map.org/). This involved drawing perpendicular lines across cortical layers using FIJI software. Proportions were calculated across three different sample areas of the entire cortex. Intensity profiles for α-synuclein and pSer129α- syn expression across cortical layers were then generated by drawing perpendicular lines and utilizing the "RGB Profile Plot" function on FIJI for the corresponding sample areas. The resulting expression values, scaled from 0 to 255, were subsequently assigned to the proportions of layers 1 to 6b obtained from the Allen Brain Atlas.

### Electron Microscopy

Brains of 12-month-old mice (n=2-3 per genotype) were fixed via intracardial perfusion with a solution of 2% PFA and 2% glutaraldehyde in 0.1M PB. This was followed by an overnight immersion in 0.1M cacodylate buffer containing 2.5% glutaraldehyde and 2% PFA^29^. The cortical layers 5 and 6 were then dissected and processed at the Yale Center for Cellular and Molecular Imaging’s Electron Microscopy Facility. Electron microscopy imaging was conducted using an FEI Tecnai G2 Spirit BioTwin Electron Microscope, and the resulting micrographs were analyzed for excitatory asymmetric synapses, their synaptic vesicles and clathrin coated vesicles using FIJI software, with the analysis performed blind to genotype.

### Nuclei isolation from cerebral cortex

Fresh cortical tissue were dissected from left hemisphere of 12 month old WT, Gba, SNCA tg and Gba-SNCA mice after euthanasia. Single nuclei were isolated as previously described with modifications^31^. All procedures were carried out on ice or at 4°C. Briefly, fresh cortical tissue was homogenized in 8.4 ml of ice-cold nuclei homogenization buffer [2 M sucrose, 10 mM Hepes (pH 7.5), 25 mM KCl, 10% glycerol, 1 mM EDTA (pH 8.0), and ribonuclease (RNase) inhibitors freshly added (40U/ml)] using a Wheaton Dounce tissue grinder (10 strokes with the loose pestle and 10 strokes with the tight pestle). The homogenate was carefully transferred into a 15 ml ultracentrifuge tube on top of 5.6 ml of fresh nuclei homogenization buffer cushion and centrifuged at 25,000 rpm for 60 min at 4°C in an ultracentrifuge. The supernatant was removed, and the pellet was resuspended in 1 ml of nuclei resuspension buffer [15 mM Hepes (pH 7.5), 15 mM NaCl, 60 mM KCl, 2 mM MgCl2, 3 mM CaCl2, and RNase inhibitors freshly added (40U/ml)] and counted on a hemocytometer with Trypan Blue staining. The nuclei were centrifuged at 500g for 10 min at 4°C with a swing bucket adaptor. They were subsequently resuspended at a concentration of 700 to 1200 nuclei/μl in the nuclei resuspension buffer for the next step of 10x Genomics Chromium loading and sequencing.

### Droplet-based single nucleus RNA sequencing and data alignment

After quality control, we recovered a total of 104,750 nuclei, including 31,906 nuclei from WT, 28,568 from Gba mutant, 26,579 from SNCA tg, and 17,697 from Gba-SNCA cortices. The mean reads per nuclei was 34,312 and the median number of identified genes per nuclei was 2410 in all samples. The snRNA-seq libraries were prepared by the Chromium Single Cell 3′ Reagent Kit v3.1 chemistry according to the manufacturer’s instructions (10x Genomics). The generated snRNA-seq libraries were sequenced using Illumina NovaSeq6000 S4 at a sequencing depth of 300 million reads per sample. For snRNA-seq of brain tissues, a custom pre-mRNA genome reference was generated with mouse genome reference (available from 10x Genomics) that included pre-mRNA sequences, and snRNA-seq data were aligned to this pre-mRNA reference to map both unspliced pre-mRNA and mature mRNA using CellRanger version 3.1.0. The raw data are available on NCBI GEO GSE283187 (https://www.ncbi.nlm.nih.gov/geo/query/acc.cgi?acc=GSE283187).

### Single-cell quality control, clustering, and cell type annotation

After quality control filtering by eliminating nuclei with less than 200 genes or more than 5% mitochondrial gene expression (poor quality nuclei) or more than 6,000 genes (potential doublets) per nucleus, we profiled 104,750 brain nuclei. Seurat (version 4.2.0) single cell analysis R package was used for processing the snRNA-seq data. The top 2000 most variable genes across all nuclei in each sample were identified, followed by the integration and expression scaling of all samples and dimensionality reduction using principal components analysis (PCA). Uniform Manifold Approximation and Projection for Dimension Reduction (UMAP) was then applied to visualize all cell clusters, and the classification and annotation of distinct cell types were based on known marker genes of each major brain cell type and the entire single nucleus gene expression matrix were investigated but were not used in downstream analyses.

### Differential expression (DE) analysis

Differential expression analysis for snRNA-seq data was performed using the Wilcoxon Rank Sum test using the function FindMarkers of the Seurat package (4.1.0) in R. For all cells, the threshold for differentially expressed genes (DEGs) was set as the expression log2 fold change of Mutant/WT mice being greater than 0.2 and significantly changed (p <0.05) after Bonferroni (BF)-correction for multiple comparisons and adjustment for possible confounders, using default parameters. Adjustment was made for differences in sex, batch and read depth with MAST^77^.

### Gene ontology (GO) pathway and targeted gene expression analysis

Gene-set and protein enrichment analysis was performed using the function enrichGO from the R package clusterProfiler in Bioconductor (3.14)^78^, with the DEGs that were significant after correction (see above) as input. Cnet plots were produced using the same package. The top three GO terms from biological process (BP) based on the lowest p-value were identified and plotted in a heatmap, without selection. Molecular function (MF) subontologies were shown instead of BP in a few cases (< 8) where BP contained duplicates or triplicates of the same pathway, for illustrative purposes, with the same DEGs and ranking used for BP and MF subontologies. The background genes were set to be all the protein-coding genes for the mouse reference genome. Default values were used for all parameters. In GO pathway analysis DEGs were entered only if significant after genome-wide BF correction (for the total number of expressed genes), using Seurat default settings, and in targeted analyses of genes of interest – i.e., genes known to function in the synapse as defined by the SynGO consortium (https://syngoportal.org)- after BF correction for the number of investigated genes. For the latter analysis, 1203 genes were found expressed in the dataset.

### Statistics

For motor behavioral studies, two-way repeated measure ANOVA followed by Sidak’s multiple comparison test was used. For cognitive behavior assays, one-way ANOVA followed by Sidak’s multiple comparison tests or Student’s t-test with Welch’s correction was used. For immunohistochemistry experiments, one-way ANOVA followed by Sidak’s multiple comparison tests was used.

## Acknowledgement

This research was funded by National Institute of Health (NIH-NINDS, 1RF1NS110354-01) R01 grant, Bell Research Initiative for Patients with Parkinson’s Disease, and Parkinson’s Foundation Research Center of Excellence (PF-RCE-1946) grant to S.S.C, US Department of Defense Early Investigator Research award (W81XWH-19-1-0264) to D.J.V., and in part by the Michael J. Fox Foundation for Parkinson’s Research (MJFF-020160) to S.S.C. and D.J.V. D.J.V.. We thank the Yale Neuroscience Rodent Behavior Analysis facility and Yale Neuroscience Imaging Core supported by The Kavli Institute of Neuroscience for usage of their microscopes. We thank Betül Yücel for contributing to genotyping of SNCA tg mice.

## Author Contribution

D.J.V., D.B., and S.S.C. conceptualized the study. D.J.V. conducted behavioral experiments and analyzed the data, prepared samples for histology and Western blotting, and performed immunohistochemistry, imaging, and image analyses. D.B. prepared samples for snRNA-seq and analyzed the snRNA-seq data. R.C. analyzed behavioral data and performed immunohistochemistry, Western blotting, imaging, and image analyses. J.R. set up founder breeding colonies and conducted genotyping for Gba L444P/KO and Gba-SNCA mice. J.P. assisted with snRNA-seq data analysis. P.M. contributed to the initial conceptualization of the study and provided founder colonies of Gba L444P/KO mice. D.J.V., D.B., and S.S.C. wrote the manuscript. All authors have read and contributed to the manuscript.

