## Supplementary Figures for "Synaptic vesicle endocytosis deficits underlie GBA-linked cognitive dysfunction in Parkinson’s disease and Dementia with Lewy bodies"

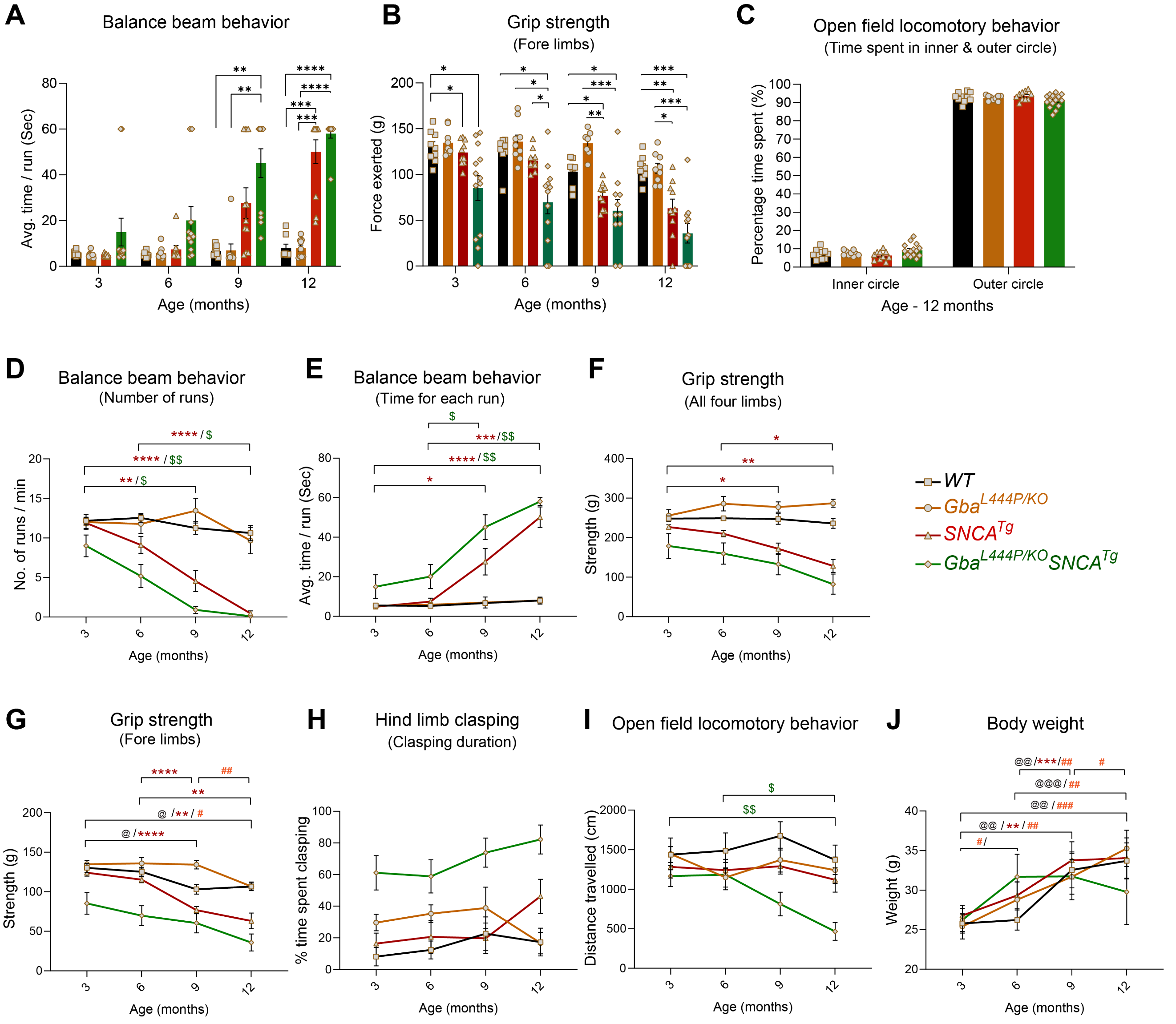


**Supplementary Figure 1. Age-related progressive worsening of motor behavior deficits in SNCA tg and Gba-SNCA mice.** Longitudinal motor behavior evaluations for WT, Gba, SNCA tg, and Gba-SNCA double mutant mice across 3, 6, 9 and 12 months of age. **A.** Balance beam behavior test showing time taken per each run. **B.** Grip strength of forelimbs. **C.** Time spent in inner and outer circle of an open field as a measure of anxiety-like behavior at 12 months of age. **D.** Balance beam behavior test showing age-related changes in number of runs per minute. **E**. Balance beam behavior test showing age-related changes in time taken per each run. **F**. Age related changes in grip strength (all four limbs). **G**. Age related changes in forelimb grip strength. **H**. Age related changes in hind limb clasping. **I**. Age related changes in open field behavior. **J**. Age related changes in body weight. n = 9-12 mice, sex-balanced cohorts. Data are presented as mean ± SEM. @ represent WT, # represent Gba mice, * represent SNCA tg mice, and $ represent Gba-SNCA double mutant mice. $^{@}p < 0.05$, $^{@@}p < 0.01$, $^{@@@}p < 0.001$, $^{@@@@}p < 0.0001$, $^{\#}p<0.05$, $^{\#\#}p < 0.01$, $^{\#\#\#}p < 0.001$, $^{\#\#\#\#}p < 0.0001$, $^{*}p < 0.05$, $^{**}p < 0.01$, $^{***}p < 0.001$, $^{****}p < 0.0001$ and, $^{\$}p < 0.05$, $^{\$\$}p < 0.01$.

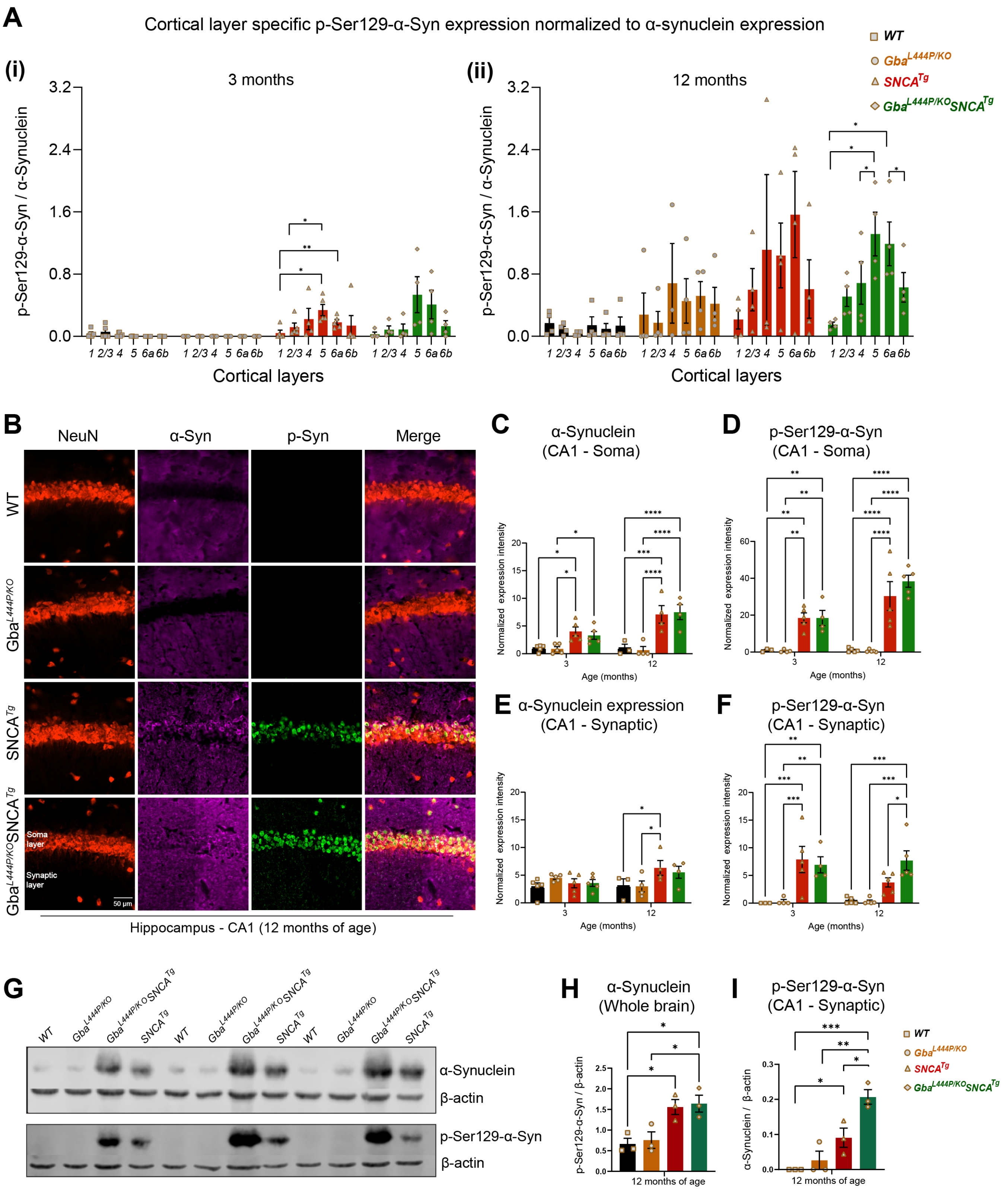


**Supplementary Figure 2. α-Synuclein and pSer129-α-syn expression in cortex, hippocampus, and whole brain. A.** Cortical layer specific pSer129-α-syn expression normalized to α-synuclein expression at the respective ages of (i) 3 and (ii) 12 months. **B**. Representative images showing α-synuclein and pSer129-α-syn expression in the CA1 subregion of the hippocampus in 12 month old mice. Scale = 50 µm. **C.** Quantification of α-synuclein expression in the soma layer of CA1 hippocampus. **D.** Quantification of pSer129-α-syn expression in the soma layer of CA1 hippocampus. **E**. Quantification of α-synuclein expression in the synaptic layers of CA1 hippocampus. **F**. Quantification of pSer129-α-syn expression in the synaptic layers of CA1 hippocampus. **G.** Western blot showing α-synuclein and pSer129-α-syn expression in the whole-brain homogenates of WT, Gba mutant, SNCA tg, and Gba-SNCA tg mice at 12 months of age. **H.** Quantification of western blots for α-synuclein levels in whole brain. **I**. Quantifications of western blots for pSer129-α-syn levels in the whole brain. n = 3 -6 mice, equal number of males and females were used. Data are presented as mean ± SEM. $*p < 0.05$, $**p < 0.01$, $***p < 0.001$, and $****p < 0.0001$.

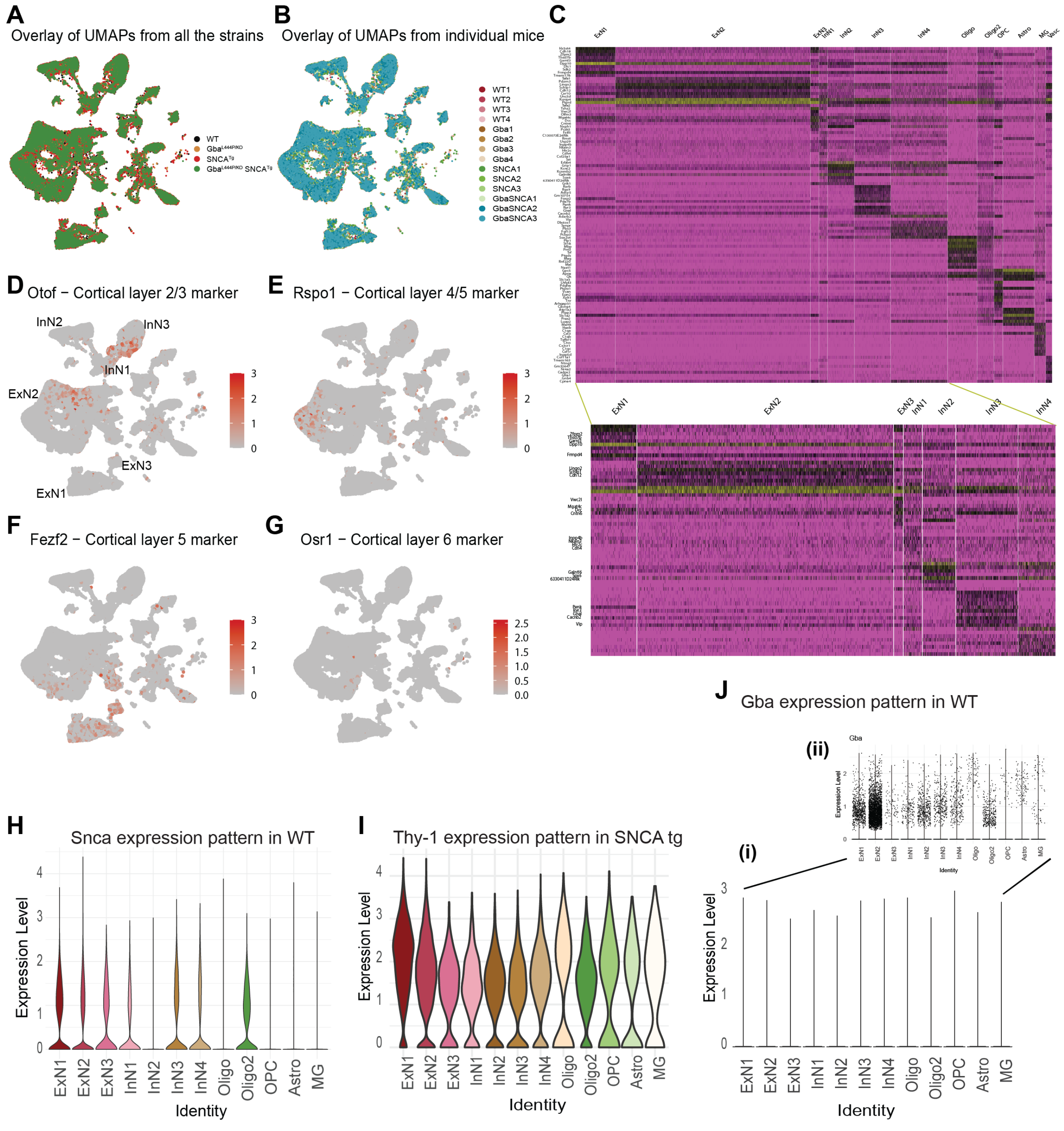


**Supplementary Figure 3. UMAPs and expression of marker genes. A.** Overlay of UMAPs for WT, Gba, SNCA tg, and Gba-SNCA showing lack of large-scale differences due to genotype. **B.** UMAPs for individual WT, Gba, SNCA tg, Gba-SNCA cortical samples. **C.** Cell type markers for the major clusters. **.** Cortical layer specific markers overlayed over UMAPs. Note, that Fezf2, the marker for Layer 5 neurons, labels ExN1 neuronal subcluster. **H.** Violin plots for expression of Snca in WT for denoted cell types. **I.** Violin plots for expression of Thy-1 in SNCA tg for denoted cell types. **J**. Violin (i) and dot (ii) plots for expression of Gba in WT for denoted cell types.

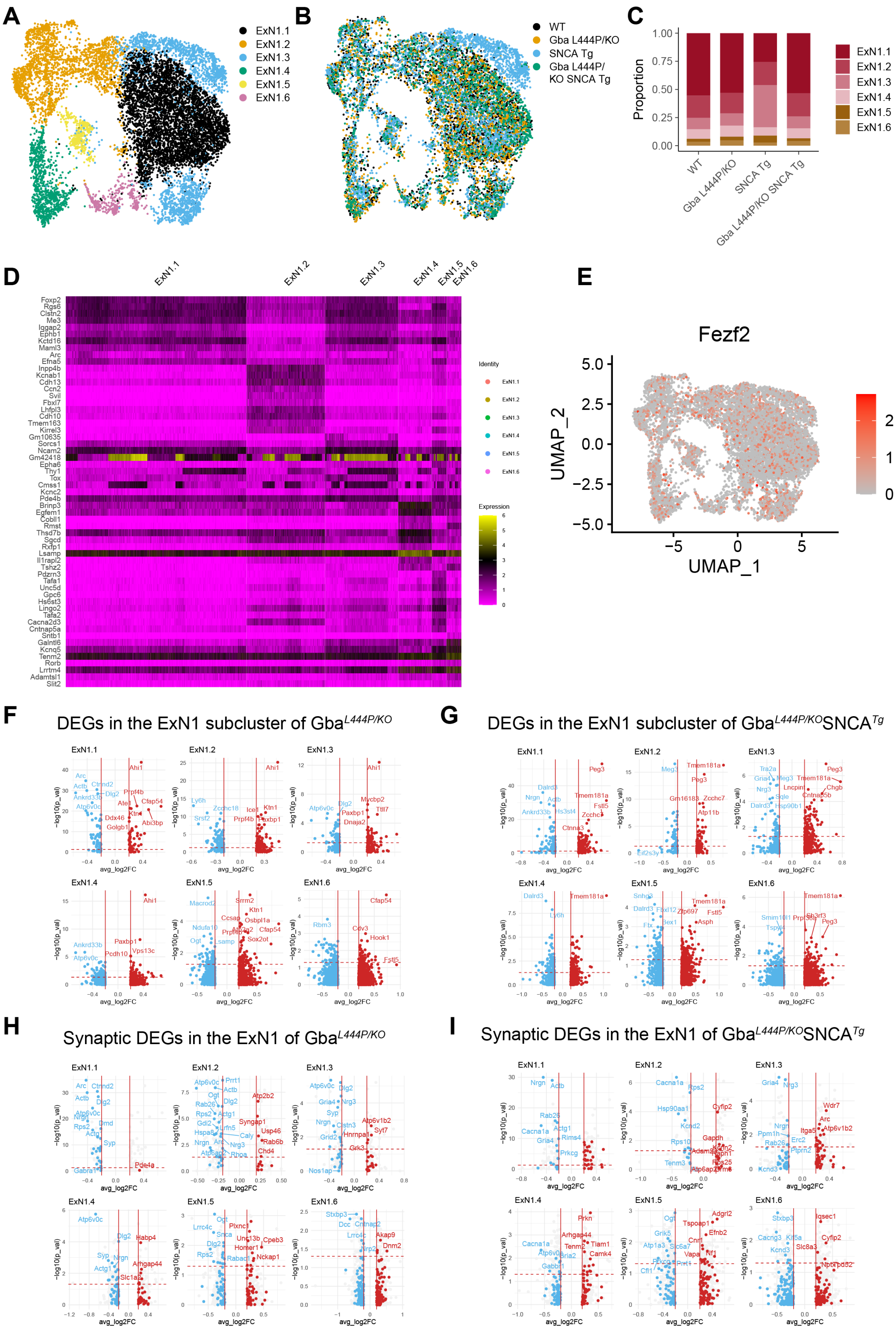

A
ExN1.1
ExN1.2
ExN1.3
ExN1.4
ExN1.5
ExN1.6
B
WT
Gba L444P/KO
SNCA Tg
Gba L444P/ KO SNCA Tg
C
Proportion
1.00
0.75
0.50
0.25
0.00
WT
Gba L444P/KO
SNCA Tg
Gba L444P/KO SNCA Tg
ExN1.1
ExN1.2
ExN1.3
ExN1.4
ExN1.5
ExN1.6
D
ExN1.1
ExN1.2
ExN1.3
ExN1.4
ExN1.5
ExN1.6
Foxp2
Rgs6
Clstn2
Me3
Iqgap2
Ephb1
Kctd16
Maml3
Arc
Efna5
Inpp4b
Kcnab1
Cdh13
Ccn2
Svil
Fbxl7
Lhfpl3
Cdh10
Tmem163
Kirrel3
Gm10635
Sorcs1
Ncam2
Gm42418
Epha6
Thy1
Tox
Cmss1
Kcnc2
Pde4b
Brinp3
Egfem1
Cobll1
Rmst
Thsd7b
Sgcd
Rxfp1
Lsamp
Il1rapl2
Tshz2
Pdzrn3
Tafa1
Unc5d
Gpc6
Hs6st3
Lingo2
Tafa2
Cacna2d3
Cntnap5a
Sntb1
Galntl6
Kcnq5
Tenm2
Rorb
Lrrtm4
Adamtsl1
Slit2
Identity
Expression
E
Fezf2
UMAP_2
UMAP_1
F
DEGs in the ExN1 subcluster of Gba^L444P/KO
G
DEGs in the ExN1 subcluster of Gba^L444P/KO SNCA^Tg
H
Synaptic DEGs in the ExN1 of Gba^L444P/KO
I
Synaptic DEGs in the ExN1 of Gba^L444P/KO SNCA^Tg
−log10(p_val)
avg_log2FC

**Supplementary Figure 4: Cortical transcriptomic signatures of ExN1. A**. UMAP showing the six subclusters found in ExN1. **B**. Overlay of the six subclusters of ExN1 in WT, Gba, SNCA tg, and Gba-SNCA samples. **C**. Proportion of cells in each ExN subcluster in the four genotypes. **D**, Heatmap showing characteristic cell type marker expression in the subclusters of ExN1. Note that high Arc expression is a characterizing marker trait of the largest subcluster ExN1.1. **E.** UMAP showing layer 5 marker Fezf2 expression pattern across ExN1 subclusters. **F.** DEGs found in the subclusters of ExN1 in Gba mice. **G.** DEGs found in the subclusters of ExN1 in Gba-SNCA mice. **H.** SynGO analysis showing synapse associated DEGs in ExN1 subclusters of Gba cortex. **I.** SynGO analysis showing synapse associated DEGs in ExN1 subclusters of Gba-SNCA cortex. Note that among all DEGs in ExN1 (**F** and **G**), DEGs are evenly distributed within up and down regulated DEGs, while the synapse associated DEGs (**H** and **I**) are selectively downregulated in all Gba as well as Gba-SNCA ExN1 subclusters. The downregulated DEGs involved in synapse vesicle cycle pathways were largely the same as shown in the main analyses, covering all cortical clusters in Fig. 4A-B and E-F, but in addition shows *Rab26* to be similarly downregulated in both genotypes (in ExN1.2 and ExN1.1, respectively).

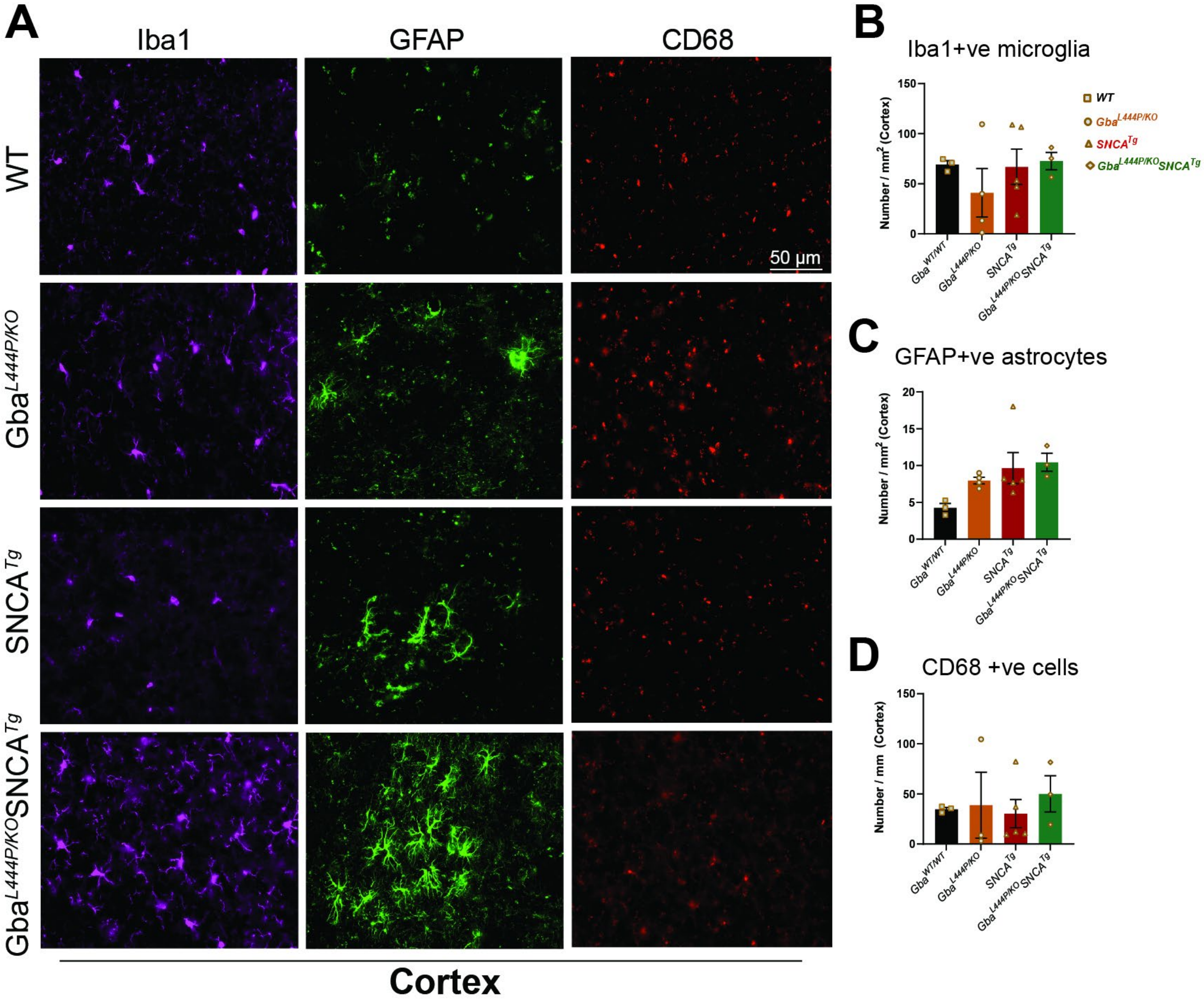


**Supplementary Figure 5. Glial signatures in Gba and Gba-SNCA mice. A**. Representative images showing cortical expression of Iba1, GFAP, and CD68 in WT, Gba, SNCA tg, and Gba-SNCA tg mice at 12 months of age. **B**. Number of Iba1 positive microglia in the cortex at 12 months. **C**. Number GFAP positive astrocytes in the cortex at 12 months. **D.** Number of CD68 (activated microglia marker) positive cells in the cortex at 12 months. Data are presented as mean ± SEM. Scale = 50 µm. * $p<0.05$, ** $p<0.01$. n=4-5 brains/genotype.

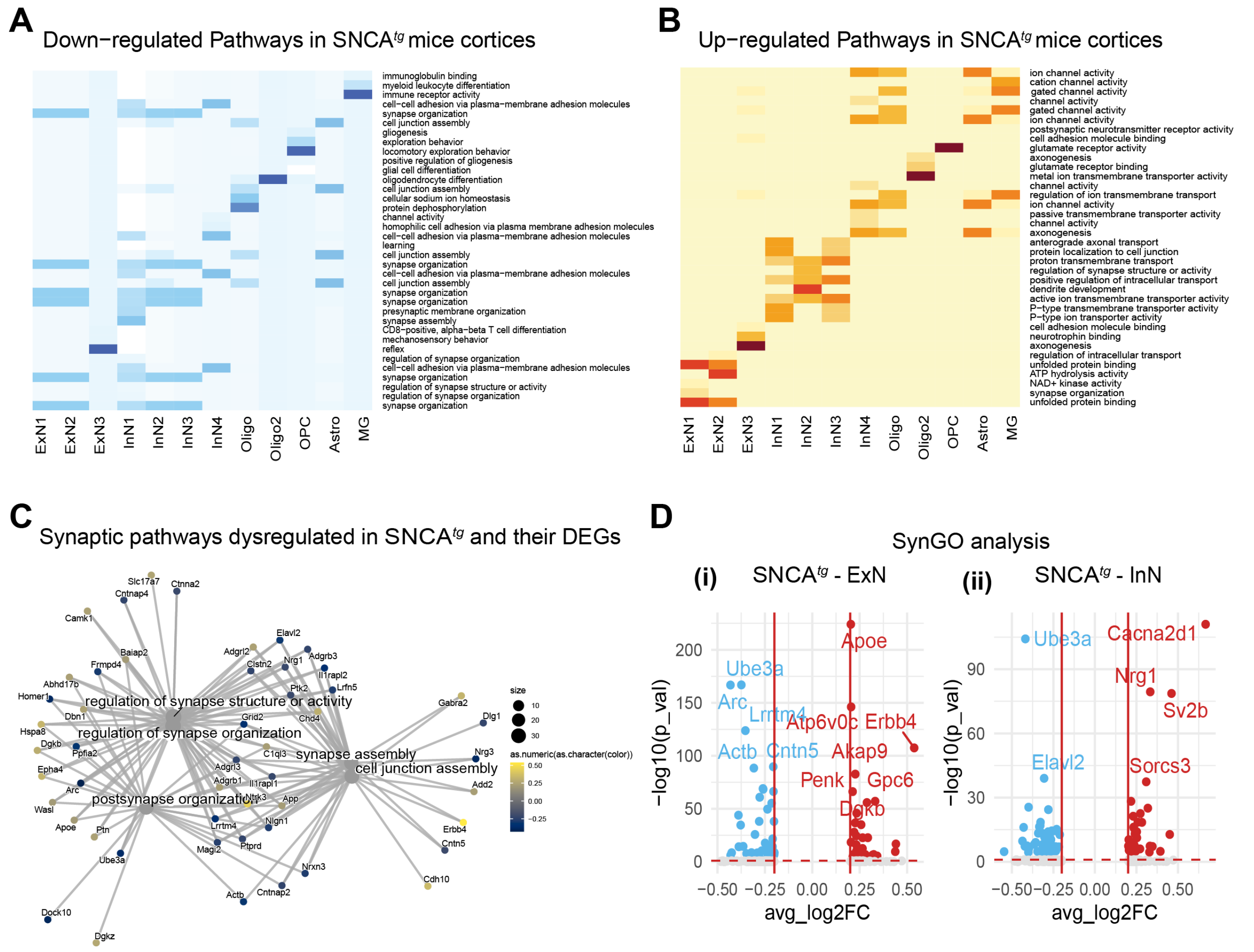


**Supplementary Figure 6. Transcriptional signatures of SNCA tg cortex**. **A-B**, Heatmap showing the most significant cortical gene ontology (GO) biological pathway alterations for each cell type, in 12-month old SNCA tg mice as revealed by unbiased analysis of enrichment of genome-wide corrected DEGs.. **C**. Cnet plot for synapse pathways with associated DEGs in SNCA tg mice **D**. SynGO analysis of DEGs in ExN and InNs of SNCA tg mice.
